# NAT10 governs uterine function and fertility by stabilizing progesterone receptor mRNA via ac^4^C modification

**DOI:** 10.1101/2025.05.27.656376

**Authors:** Shi Tang, Da-Ke Peng, Run-Fu Jiang, Yu-Qi Hong, Qing-Yan Zhang, Xiao-Qi Yang, An-Nan Zhang, Yan-Wen Xu, Shi-Hua Yang, Ji-Long Liu

## Abstract

The *N*^4^-acetylcytidine (ac^4^C) modification is one of the most abundant chemical modifications in mammalian transcriptome, which plays a crucial role in regulating gene expression across various biological processes. In this study, we investigate the role of *N*^4^-acetyltransferase 10 (NAT10), the sole enzyme responsible for catalyzing the ac^4^C modification, in uterine function. Our findings revealed that knockdown of NAT10 in cultured human endometrial stromal cells leads to decreased progesterone receptor (PGR) expression and compromised decidualization. Mechanistically, NAT10 is found to enhance the stability of *PGR* mRNA through its ac^4^C modification in the coding sequence (CDS). Furthermore, conditional deletion of *Nat10* in the mouse uterus using *Pgr*-Cre resulted in defective pubertal uterine development, characterized by a thinner stroma and reduced endometrial gland number. These mice were infertile in adulthood, experiencing failures in both implantation and decidualization. However, heterozygous *Nat10* conditional knockout mice showed reduced NAT10 expression in the uterus without affecting uterine development or fertility. To bridge the gap between heterozygous and homozygous knockout conditions, we employed Remodelin, an inhibitor of NAT10, in adult female mice, to mimic an intermediate gene dosage. Our results demonstrated that inhibiting NAT10 with Remodelin preserved uterine structures but disrupted the PGR signaling pathway, leading to impaired uterine function during pregnancy. In conclusion, our study provides compelling evidence that NAT10 safeguards uterine function and fertility by regulating the ac^4^C modification of PGR mRNA.

**Significance:** Here we identified NAT10-catalyzed ac^4^C modification of progesterone receptor (PGR) mRNA as a crucial mechanism regulating PGR expression by enhancing its mRNA stability. In vitro studies using human endometrial stromal cells show that the NAT10-PGR axis is required for decidualization. Additionally, genetic and pharmacologic manipulations in mice further demonstrate the necessity of NAT10-dependent P_4_/PGR signaling for uterine function during implantation and decidualization. Our findings highlight the importance of NAT10-mediated ac^4^C modification in maintaining PGR expression and uterine function.

## Introduction

The uterus is an essential organ for mammalian gestation. At birth, both the endometrial and myometrial components of the uterus are immature, with the primary architecture forming before puberty [1]. During puberty, under the influence of ovarian 17β-estradiol (E_2_) and progesterone (P_4_), the uterus grows and acquires optimal responsiveness via estrogen receptor alpha (ESR1) and progesterone receptor (PGR). After puberty, the endometrial epithelial and stromal cells follow a cyclic pattern of proliferation and differentiation in response to E_2_ and P_4_, preparing the uterus for potential embryo implantation within a ∼28-day menstrual cycle: in the proliferative phase, E_2_ stimulates epithelial proliferation, while P_4_ sustains stromal proliferation and drives decidualization in the secretory phase [2, 3]. Similar hormonal-driven changes also occur in mice, albeit in the context of pregnancy, as functional corpus luteum development in mice is mating/fertilization-dependent [4]. In early gestation in mice, E_2_ stimulates the proliferation of uterine epithelial cells on gestational days (GD) 1-2. Subsequently, P_4_ sustains stromal proliferation while inhibiting epithelial proliferation, thereby preparing the uterus on GD4 for embryo attachment. Upon attachment of the embryo to the uterine epithelium, the stromal cells located beneath undergo a cessation of proliferation and differentiate into decidual cells, which play a pivotal role in facilitating trophoblast invasion and placentation [5]. Over the past few decades, extensive research has been conducted on the components of E_2_ and P_4_ signaling essential for uterine development and function [6]. However, a comprehensive molecular map of these signaling pathways remains elusive.

In recent years, emerging evidence underscores the pivotal regulatory role of posttranscriptional mRNA modifications, collectively known as the epitranscriptome, in the dynamics of gene expression. A diverse array of chemical modifications occurs on mRNA, including *N*^6^-methyladenosine (m^6^A), 5-methylcytosine (m^5^C), *N*^4^-acetylcytosine (ac^4^C), *N*^7^-methylguanosine (m^7^G), and *N*^1^-methyladenosine (m^1^A). Prior research has predominantly centered on m^6^A and m^5^C modifications, whereas studies on ac^4^C modification remain relatively scarce [7]. Initially, ac^4^C was identified in tRNA and rRNA [8]. However, with the advancement of analytical chemistry and high-throughput sequencing technologies, recent discoveries have unveiled the widespread presence of ac^4^C modifications on mRNA. In fact, ac^4^C ranks as the third most abundant modification in the transcriptome [9]. Notably, it stands as the sole acetylation event observed in eukaryotic mRNA. This modification is predominantly enriched within the coding sequence (CDS) of mRNA, serving to enhance mRNA stability and translation efficiency [10]. The ac^4^C modification is catalyzed by the enzyme *N*-acetyltransferase 10 (NAT10), which is the only human enzyme known to possess both acetyltransferase and RNA-binding activities [11]. Accumulating evidence suggests that ac^4^C is indispensable for a multitude of biological processes, including spermatogenesis, oogenesis, and embryonic development [12]. However, the precise role of mRNA ac^4^C in uterine function and pregnancy remains unknown.

Here, we identified PGR as a direct target of NAT10-mediated ac^4^C modification. Using cultured human endometrial stromal cells (hESCs), we demonstrated that the NAT10-PGR axis is required for in vitro decidualization. Using a combination of genetic and pharmacologic tools in mouse models, we demonstrated that NAT10-dependent P_4_/PGR signaling is necessary to maintain uterine function during implantation and decidualization. In summary, our results provide evidence that NAT10 safeguards uterine function and fertility by regulating ac^4^C modification of *PGR* mRNA.

## Results

### NAT10 expression in hESCs is required for decidualization in vitro

To gain insights into the role of NAT10 in normal endometrial physiology, we initially utilized immunochemistry to analyze NAT10 expression in endometrial biopsy samples collected from healthy volunteers with regular menstrual cycles. Consistent with results from a microarray dataset [13] (**Fig. S1**), our findings revealed that the NAT10 protein was abundantly present in both epithelial and stromal cells during the proliferative phase. However, its expression decreased during the early secretory phase. Notably, NAT10 expression rebounded in the middle secretory phase but subsequently declined again in the late secretory phase (**Fig. 1A**). This fluctuating pattern of expression suggests that NAT10 may play a crucial role during the peri-implantation period. We subsequently explored whether dysregulation of NAT10 is linked to human implantation disorders. Previous studies have employed single-cell RNA-seq to analyze the endometrium of patients with recurrent implantation failure (RIF) during the mid-secretory phase [14] and the fetal-maternal interface of patients with recurrent pregnancy loss (RPL) in the first trimester [15]. We generated pseudo-bulk RNA-seq data for epithelial and stromal cells by averaging single-cell RNA-seq data, following the methodology described in our earlier study [16, 17]. Our results showed no significant change in NAT10 expression in either endometrial epithelial cells or stromal cells from RIF patients compared to fertile controls (**Fig. 1B**). In contrast, we observed a significant downregulation of NAT10 expression in stromal cells from RPL patients compared to fertile controls (**Fig. 1C**). Furthermore, we found that the downregulated genes in stromal cells from RPL patients were significantly enriched for decidualization-related genes, including PRL, PGR, and HOXA11 (**Fig. S2A-C**). These findings led us to speculate that the expression of NAT10 in stromal cells is associated with the process of decidualization.

**Figure 1.**
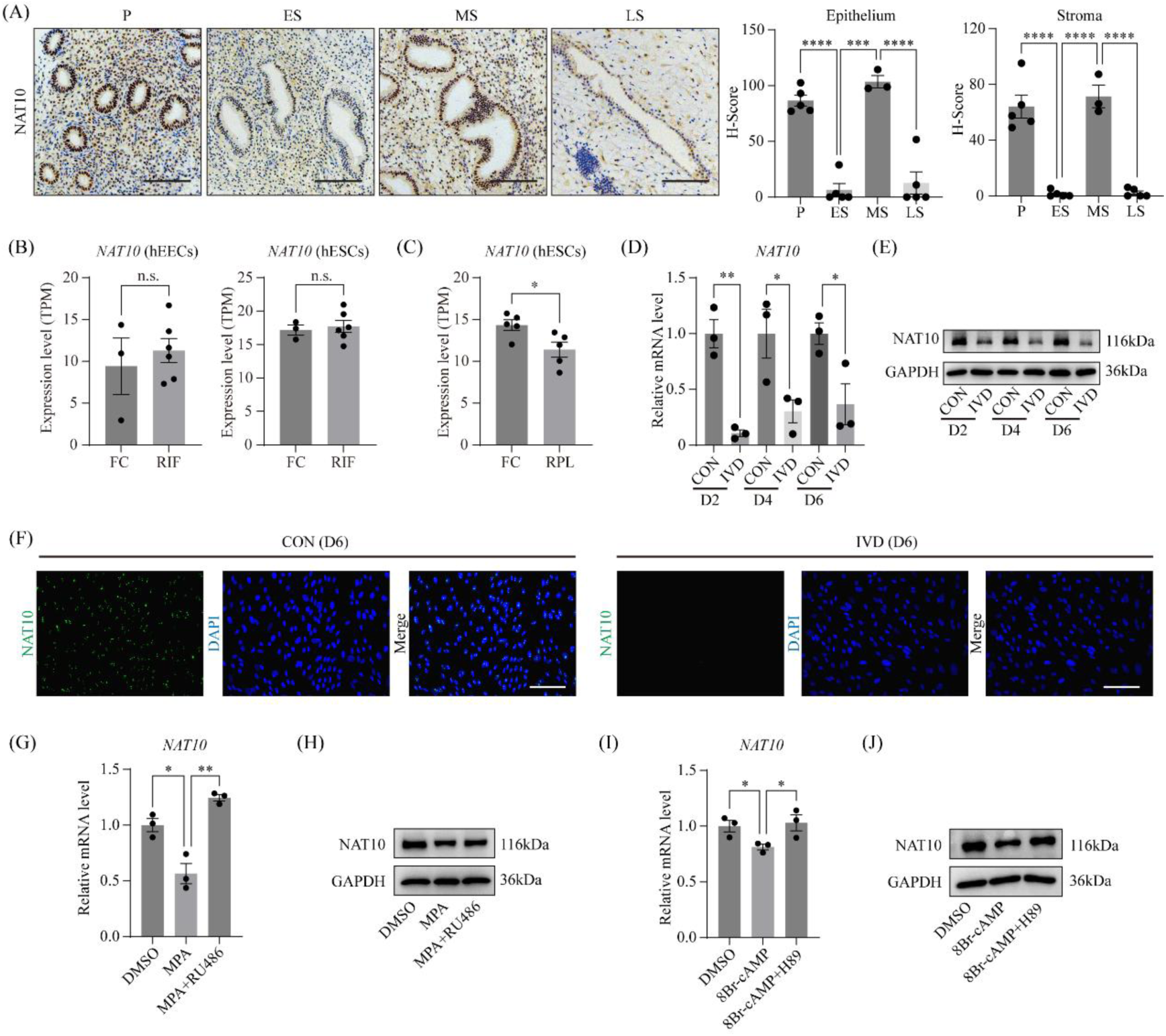
Dynamic expression of NAT10 in human endometrial stromal cells during decidualization. (A) Immunohistochemical analysis reveals the expression pattern of NAT10 protein in the endometrium throughout the menstrual cycle. P, proliferative phase; ES, early secretory phase; MS, middle secretory phase; LS, late secretory phase. Scale bar =100 μm. (B) The mRNA expression levels of *NAT10* in human endometrial epithelial cells (hEECs) and endometrial stromal cells (hESCs) from fertile control patients (FC) and patients with recurrent implantation failure (RIF), as obtained from the single-cell RNA-seq dataset GSE183837. Data are presented as means ± SEM. n.s., not significant. (C) The mRNA expression levels of *NAT10* in hESCs from FC patients and patients with recurrent pregnancy loss (RPL), based on the single-cell RNA-seq dataset GSE174399. Data are presented as means ± SEM. *, P < 0.05. (D) Quantitative RT-PCR analysis of *NAT10* RNA expression levels in decidualizing hESCs in vitro at various time points. CON, vehicle control; IVD, in vitro decidualization. Data are presented as mean ± SEM. *, P < 0.05; **, P < 0.01. (E-F) Protein expression levels of NAT10 in hESCs during decidualization, as measured by western blot (E) and immunofluorescent staining (F). (G-H) Expression of *NAT10* mRNA and protein in hESCs treated with MPA alone or MPA combined with RU486 for 2 days. Data are presented as mean ± SEM. *, P < 0.05; **, P < 0.01. (I-J) Expression of *NAT10* mRNA and protein in hESCs treated with 8Br-cAMP alone or 8Br-cAMP combined with H89 for 2 days. Data are presented as mean ± SEM. *, P < 0.05.

To elucidate the crucial role of NAT10 in the process of decidualization, we developed an in vitro model utilizing primary human endometrial stromal cells (hESCs). These cells were induced to undergo decidualization through a combined treatment regimen of the P_4_ analog MPA and the cAMP analog 8-Br-cAMP. The success of in vitro decidualization was confirmed by the robust induction of PRL and IGFBP1 (**Fig. S3A-B**), which are well-established markers of decidualization. Our model demonstrated that the transition of hESCs into a decidualized state is accompanied by a notable reduction in NAT10 expression, evident at both the mRNA and protein levels (**Fig. 1D-F**). Intriguingly, we observed a similar decline in NAT10 levels in the uterus of pregnant mice. Specifically, NAT10 protein is abundantly expressed in stromal cells of the mouse uterus during the implantation window (GD4-5) (**Fig. S4A**). However, at GD8, the expression of NAT10 in the primary decidual zone (PDZ) was significantly lower compared to that in the secondary decidual zone (SDZ) (**Fig. S4B**). Furthermore, a previous spatiotemporal transcriptome dataset [18] revealed a gradual decrease in NAT10 expression in decidual cells located in the mesometrial region or decidua basalis (**Fig. S4C**). To further validate these observations, we isolated primary mouse endometrial stromal cells (mESCs) from wild-type mice on GD4 and induced them to undergo decidualization in vitro by treating them with E_2_ and P_4_. Remarkably, we found that the expression of NAT10 was significantly downregulated in mESCs following in vitro decidualization (**Fig. S4D-E**). To pinpoint the factors responsible for modulating NAT10 expression during this process, we individually treated hESCs with MPA or 8-Br-cAMP and assessed their impact on NAT10 expression. Notably, treatment with either MPA or 8-Br-cAMP alone resulted in a significant decrease in NAT10 expression. Crucially, this reduction was reversed in the presence of the antagonists RU486 or H89, respectively (**Fig. 1G-J**). Collectively, our findings reveal that NAT10 is down-regulated in the process of decidualization, which is driven by the combined action of P_4_ and cAMP signaling.

To investigate whether NAT10 influences the functional properties of hESCs, we knocked down NAT10 expression in these cells using siRNA. The effectiveness of the NAT10 knockdown was verified through RT-PCR, western blot, and immunofluorescence analyses (**Fig. S5A-C**). We found that knockdown of NAT10 slightly reduced cell proliferation (**Fig. S5D-E**), but had no discernible effect on apoptosis (**Fig. S5F**). To determine whether NAT10 expression is required for the decidualization process, we knocked down NAT10 in hESCs prior to treating them with MPA and 8-Br-cAMP. Compared to control cells, we observed a marked decrease in the expression of PRL and IGFBP1 in NAT10-knockdown hESCs after 2, 4, and 6 days of decidualization (**Fig. 2A-B**). Typically, hESCs exhibit an elongated fibroblast-like morphology, whereas decidualized hESCs adopt a polygonal epithelioid shape. Following NAT10 knockdown, we observed a significant impairment in the decidualization capacity of hESCs, evident from the reduced cell shape transformation (**Fig. 2C**). To further validate the impact of NAT10 expression on decidualization, we knocked down NAT10 at day 2 and day 4 after initiating decidualization and found that the expression of PRL and IGFBP1 was consistently decreased at day 6 of decidualization (**Fig. 2D-E**). Given that NAT10 is downregulated during the process of decidualization, we explored whether overexpressing NAT10 affects the ability of hESCs to undergo decidualization. As anticipated, NAT10 overexpression led to decreased expression of PRL and IGFBP1 in hESCs decidualized for 4 days (**Fig. 2F-G**). These data suggest that NAT10 is essential throughout the process of decidualization, yet excessive NAT10 expression disrupts this process, indicating that the levels of NAT10 expression during decidualization must be finely tuned.

**Figure 2.**
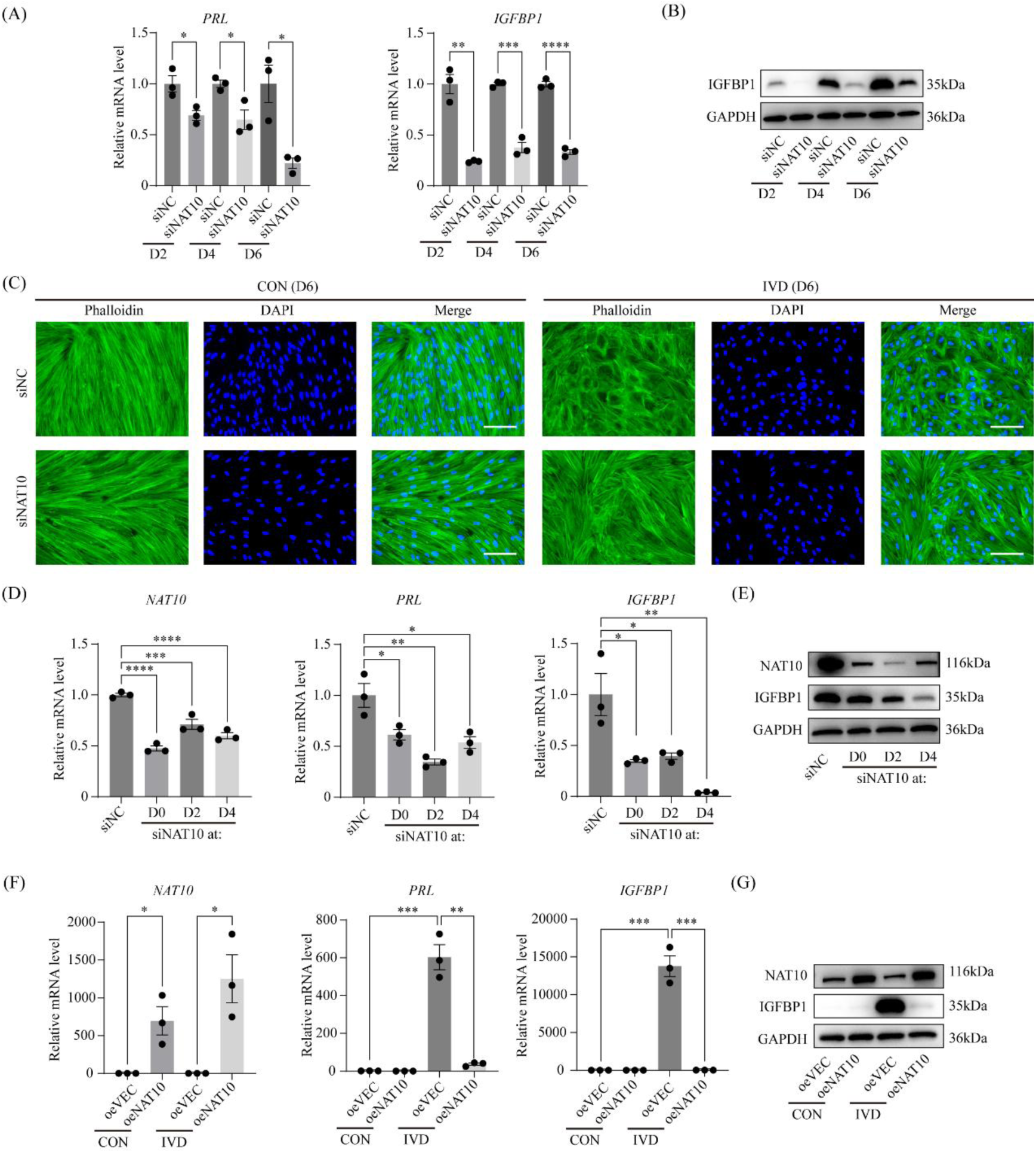
The necessity of NAT10 expression in hESCs for the process of decidualization. (A) Quantitative RT-PCR analysis of *PRL* and *IGFBP1* RNA levels in hESCs, following NAT10 knockdown and subsequent decidualization for durations of 2, 4, or 6 days. siNC, non-targeting control; siNAT10, NAT10-targeting siRNA. Data are presented as mean ± SEM. *, P < 0.05; **, P < 0.01; ***, P < 0.001; ****, P < 0.0001. (B) Protein expression levels of IGFBP1 following NAT10 knockdown and decidualization. (C) Phalloidin staining of F-actin in decidualized hESCs after siRNA-mediated NAT10 knockdown for 6 days. Bar = 100 μm. (D) RNA expression levels of *PRL* and *IGFBP1* in decidualized hESCs at day 6 when transfected with NAT10 siRNA at days 0, 2, and 4 post-decidualization induction. Data are presented as mean ± SEM. *, P < 0.05; **, P < 0.01; ***, P < 0.001; ****, P < 0.0001. (E) Protein expression levels of IGFBP1 corresponding to NAT10 siRNA transfection at days 0, 2, and 4 post-decidualization induction. (F) Quantitative RT-PCR analysis of *PRL* and *IGFBP1* mRNA expression in hESCs following NAT10 overexpression and subsequent decidualization for 4 days. oeVEC, empty vector control; oeNAT10, NAT10 overexpression vector. Data are presented as means ± SEM. *, P < 0.05; **, P < 0.01; ***, P < 0.001. (G) IGFBP1 protein expression in hESCs following NAT10 overexpression and subsequent decidualization for 4 days.

To investigate whether the role of NAT10 in decidualization is preserved in mice, we utilized siRNA to silence NAT10 expression in isolated mESCs prior to decidualization induction. Notably, our findings revealed a substantial decrease in the expression of *Prl8a2*, a marker of mouse decidualization, upon NAT10 knockdown (**Fig. S6A-B**). These results suggest that NAT10 maintains a conserved function in decidualization across both human and mouse species.

### NAT10 safeguards PGR expression in hESCs

It has been reported that NAT10 possesses catalytic activity not only towards RNA but also towards protein [19]. In our study, we observed that silencing NAT10 in hESCs led to a notable decrease in RNA acetylation, whereas no significant changes were detected in protein acetylation levels (**Fig. S7A-B**). These results suggest that RNA is the predominant substrate of NAT10 compared to proteins in hESCs.

To understand the mechanism underlying the function of NAT10 as the RNA ac^4^C writer in hESCs, we conducted acetylated RNA immunoprecipitation sequencing (acRIP-seq) on mRNAs isolated from human endometrial tissues at the middle secretory phase (pooled endometrial tissues from 3 patients). This analysis revealed 39583 unique ac^4^C peaks associated with 9811 genes (**Table S1**). Notably, the majority of these ac^4^C peaks were located within the coding sequence (CDS) region, accounting for 64.3% of the total peaks, followed by the intronic region (27.0%), 3′-UTR (4.2%), 5′-UTR (3.7%), and intergenic region (0.8%) (**Fig. 3A**). A metagene analysis further revealed a pronounced enrichment of ac^4^C peaks in proximity to both the start and stop codons, as evidenced by peak density (**Fig. 3B**), aligning with a previous report [20]. Additionally, an unbiased motif search uncovered a significant overrepresentation of repeating CXX (several obligate cytidines interspersed with two non-obligate nucleotides) motifs within ac^4^C peaks (**Fig. 3C**), corroborating earlier observations [10].

**Figure 3.**
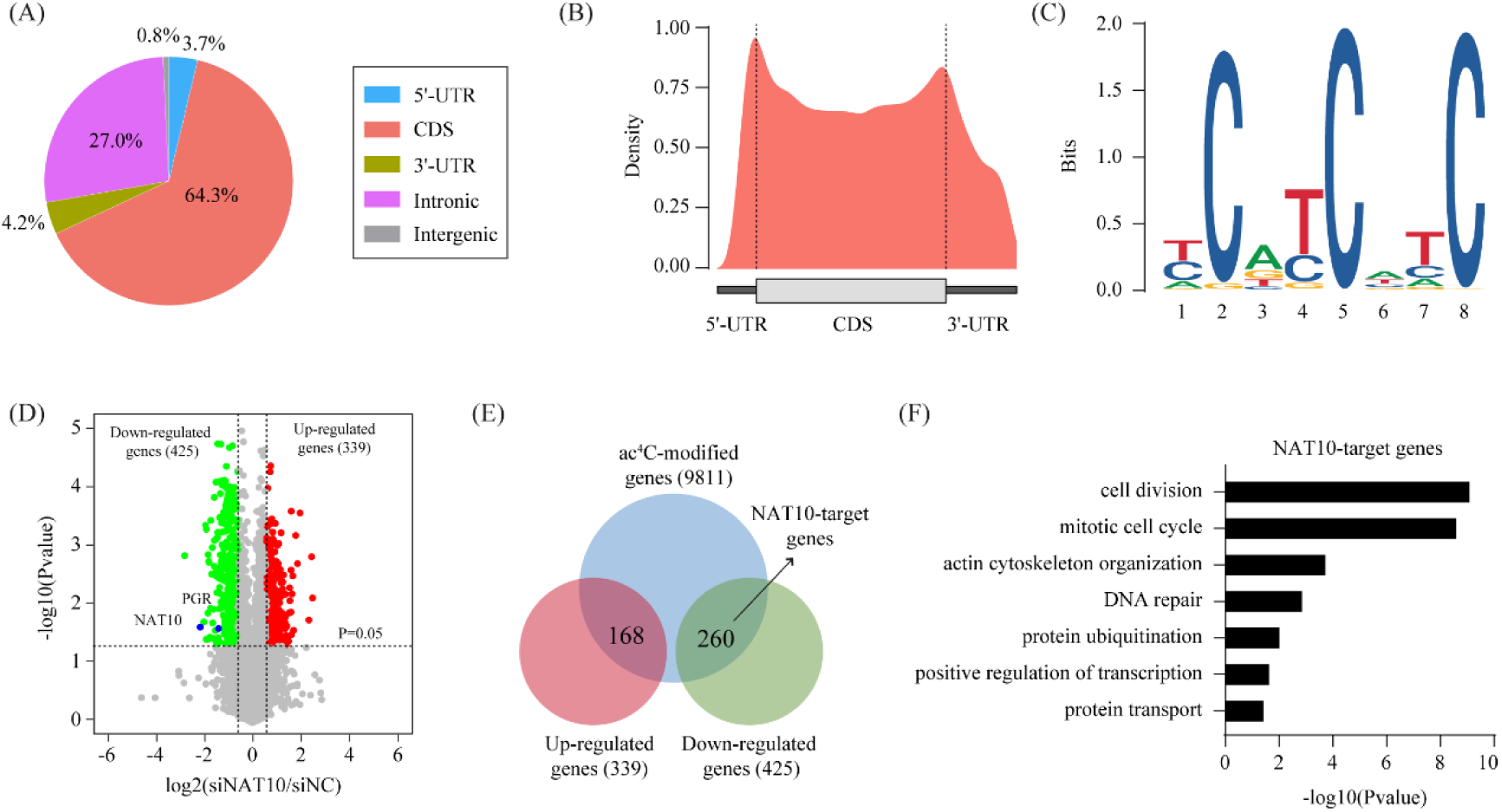
Identification of NAT10 target genes using acRIP-seq and RNA-seq. (A-C) Global profiling of ac^4^C modification in normal human endometrium via acRIP-seq. (A) Pie chart illustrating the distribution of ac^4^C peaks across various genomic segments. UTR, untranslated region; CDS, coding sequence. (B) The metagene plot showing the distribution of ac^4^C peaks within the gene body. (C) The consensus sequence motif derived from ac^4^C peaks, presented as a sequence logo. (D) Volcano plot showing significantly expressed genes in hESCs after NAT10 knockdown for 48h, as determined by RNA-seq. Differentially expressed genes were selected based on a fold change > 1.5 and a statistical significance level of P < 0.05. (E) Venn diagram depicting the overlap of differentially expressed genes identified by RNA-seq and ac^4^C-modified genes identified by acRIP-seq. (F) Gene ontology enrichment analysis of overlap genes.

To uncover genes regulated by NAT10 in hESCs, we performed RNA-seq analysis on mRNAs isolated from hESCs with and without NAT10 knockdown. Analysis of the RNA-seq data identified 764 differentially expressed genes, including 339 up-regulated genes and 425 down-regulated genes (**Fig. 3D and Table S2**). To identify direct target genes of NAT10, we integrated acRIP-seq and RNA-seq data, focusing on genes that exhibited both ac^4^C modification sites and down-regulation upon NAT10 knockdown. Using this approach, we pinpointed 260 potential direct target genes of NAT10 (**Fig. 3E**). Gene ontology (GO) analysis revealed that these genes were significantly enriched in cell division, mitotic cell cycle, actin cytoskeleton organization, DNA repair, protein ubiquitination, positive regulation of transcription, and protein transport (**Fig. 3F**).

Intriguingly, our study uncovered *PGR* as a direct target gene of NAT10. We observed an ac^4^C peak spanning the entire exon 1 for both PGR-A and PGR-B isoforms (**Fig. 4A**). An analysis of RNA-seq data further substantiated our findings, which showed a downregulation of *PGR* in hESCs with NAT10 knockdown (**Fig. 4B**). This observation was consistently supported by RT-PCR and western blot analyses, which demonstrated a significant inhibition of PGR expression upon siRNA-mediated knockdown of NAT10 (**Fig. 4C-D**) and a marked upregulation of PGR expression following NAT10 overexpression in hESCs (**Fig. 4E-F**). To ascertain whether *Pgr* serves as a conserved target of NAT10 in mice, we conducted acRIP-seq using samples from adult mouse uteri (**Table S3**). The distribution pattern of ac^4^C peaks in the mouse uterus exhibited remarkable similarity to the human acRIP-seq data (**Fig. S8A-E**). Most importantly, we identified an ac^4^C peak at the identical location of *PGR* in mice (**Fig. S8F**). Furthermore, siRNA-mediated knockdown of NAT10 in mESCs similarly resulted in a significant inhibition of PGR expression (**Fig. S8G-H**). These data suggest a conservation of ac^4^C modification on *Pgr* mRNA across both human and mouse species.

**Figure 4.**
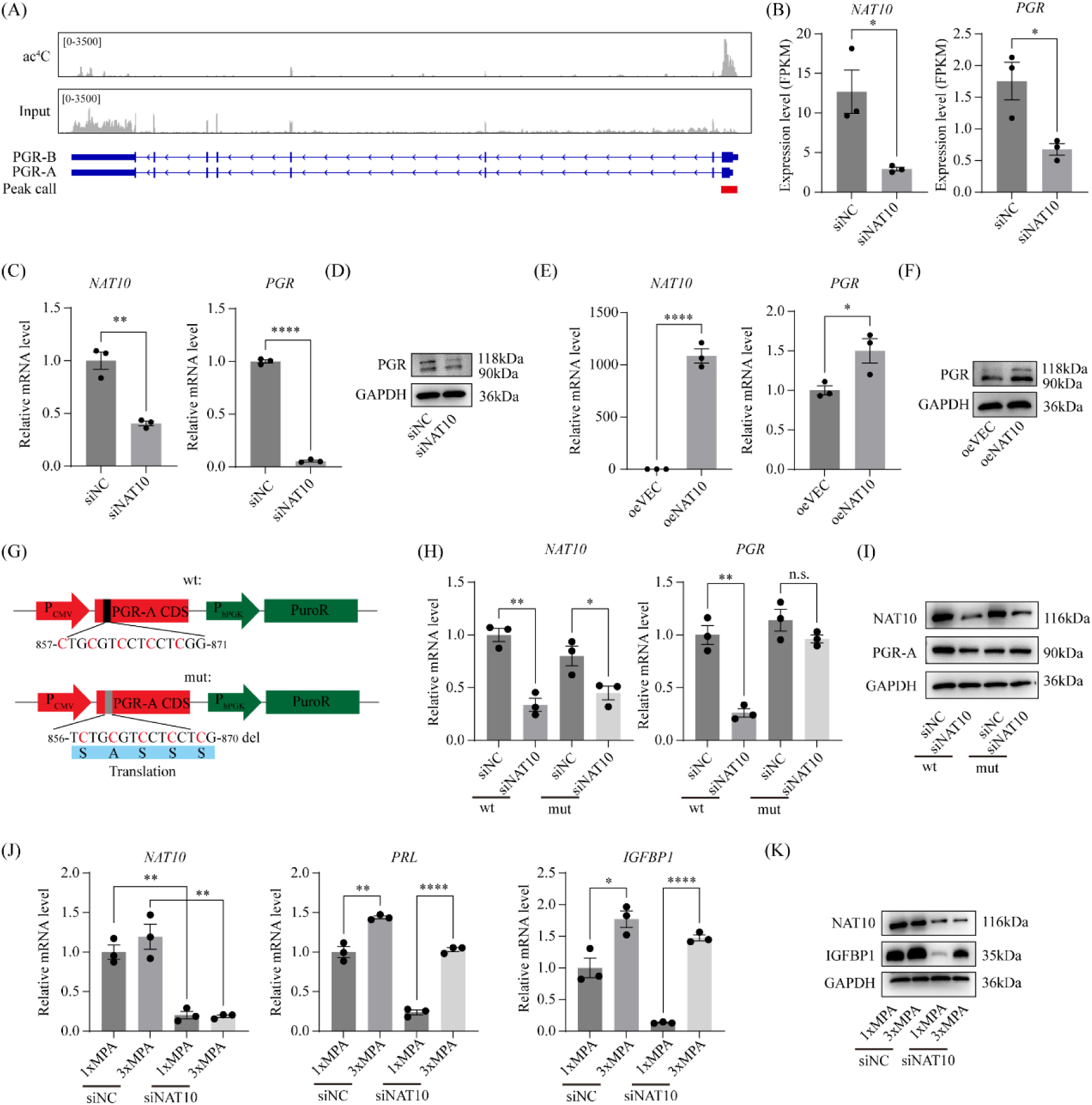
*PGR* mRNA as a direct target of NAT10-mediated ac^4^C modification. (A) Integrative Genomics Viewer (IGV) snapshot depicting the coverage of ac^4^C immunoprecipitation and input in *PGR* mRNA, based on acRIP-seq data. Peak calling was executed using MACS3 software. (B) RNA expression levels of *PGR* in hESCs following NAT10 knockdown, as derived from RNA-seq data. Data are presented as mean ± SEM. *, P < 0.05. (C-D) Validation of PGR expression levels in hESCs following 48h of NAT10 knockdown by using quantitative RT-PCR and western blot. Data are presented as mean ± SEM. **, P < 0.01; ****, P < 0.0001. (E-F) Expression of *NAT10* mRNA and protein in hESCs after 48h of NAT10 overexpression. Data are presented as mean ± SEM. *, P < 0.05; ****, P < 0.0001. (G) Schematic diagram of the reporter vector. The PGR coding sequence (CDS) was inserted downstream of the CMV promoter. The hPGK promoter-driven puromycin resistance gene (PuroR) serves as a control for transfection efficiency. A cluster of potential ac^4^C sites within the PGR CDS, which is conserved between humans and mice, was predicted using the PACES tool. (H-I) Impact of ac^4^C sites on reporter gene expression in HEK293T cells co-transfected with the reporter vector and either siNAT10 or siNC for 48 hours. Data are presented as mean ± SEM. *, P < 0.05; **, P < 0.01. (J) Quantitative RT-PCR analysis of *PRL* and *IGFBP1* mRNA levels in decidualized hESCs following NAT10 knockdown and rescue with excessive medroxyprogesterone acetate (MPA). Data are presented as means ± SEM. *, P < 0.05; **, P < 0.01; ****, P < 0.0001. (K) Western blot analysis of IGFBP1 protein expression in decidualized hESCs after NAT10 knockdown and rescue with excessive MPA.

Subsequently, our attention turned to precisely locating the ac^4^C-modified sites within the PGR mRNA. Leveraging the PACES tool [21], we discovered a potentially conserved ac^4^C site within the coding sequence (CDS) of the PGR mRNA, shared between humans and mice (**Fig. 4G**). To substantiate this finding, we conducted experiments by expressing the CDS of human PGR in HEK293T cells, both with the intact site (PGR-wt) and without it (PGR-mut). Notably, our results demonstrated that upon NAT10 knockdown, only PGR-wt, and not PGR-mut, underwent downregulation (**Fig. 4H-I**). Previous studies have shown that ac^4^C modification can enhance mRNA stability and translation efficiency [10]. However, given that the reduction in PGR expression upon NAT10 knockdown was more pronounced at the mRNA level (**Fig. 4H**) compared to the protein level (**Fig. 4I**), we speculated that the ac^4^C modification in the PGR mRNA CDS primarily functions to bolster mRNA stability rather than translation efficiency. To further investigate this, we analyzed the impact of ac^4^C on PGR mRNA stability in hESCs. By treating the cells with actinomycin-D to halt transcription, we observed a marked decrease in PGR mRNA stability after NAT10 knockdown and a marked increase in PGR mRNA stability after NAT10 overexpression (**Fig. S9A-B**). Collectively, these findings indicate that ac^4^C modification within the CDS of PGR mRNA promotes PGR expression by enhancing the stability of its mRNA.

Previous research has demonstrated that PGR protein expression peaks in the proliferative phase of the human endometrium and gradually decreases throughout the secretory phase [22]. Our study uncovered that PGR expression is downregulated in hESCs undergoing in vitro decidualization (**Fig. S10A-B**). This downregulation appeared to be mediated by MPA, as MPA alone suppressed PGR expression while cAMP alone enhanced it (**Fig. S10C-F**). Silencing PGR in hESCs hindered decidualization (**Fig. S10G-H**). Furthermore, overexpressing either PGR isoform, PGR-A or PGR-B, also impaired decidualization (**Figure S10I-L**). These observations indicate that, like NAT10, a programmed reduction in PGR level is crucial for decidualization to proceed. Consequently, our attempt to rescue NAT10 knockdown by overexpressing PGR was unsuccessful (**Fig. S11**). Alternatively, we explored the use of varying MPA concentrations to modulate PGR activity in our rescue experiments (**Fig. S12**). We discovered that supplementing with additional MPA (three times of normal concentration) was effective in restoring decidualization in hESCs with NAT10 knockdown, whereas it had minimal impact when NAT10 was intact (**Figure 4J-K**). These findings suggest that PGR functions as a pivotal downstream effector of NAT10 in hESCs during the decidualization process.

### *Pgr*-Cre-mediated deletion of NAT10 in mice results in impaired pubertal uterus development

To investigate the role of NAT10 in the mouse uterus, we generated conditional *Nat10* knockout mice (*Nat10*^d/d^) by crossing *Nat10*-floxed mice (*Nat10*^f/f^) with mice expressing Cre recombinase under the control of the *Pgr* promoter (*Pgr*^Cre/+^) (**Fig. 5A** and **Fig. S13A-C**). PCR analysis confirmed that exon 4 of *Nat10*, flanked by two floxp sites, was successfully removed by Cre recombinase in the uteri of *Nat10*^d/d^ mice at postnatal day 14 (PND14), PND21, and 6 weeks (6w) (**Fig. 5B**). RNA-seq analysis of uterine samples collected at PND21 revealed that the excision of exon 4 (172bp) induced alternative splicing between exons 3 and 5, resulting in a frameshift in the open reading frame (**Fig. S14A**). Intriguingly, RNA-seq data indicated a significant downregulation of *Nat10* mRNA upon exon 4 deletion (**Fig. S14B**), potentially due to nonsense-mediated mRNA decay (NMD). This downregulation was further validated by quantitative RT-PCR (**Fig. S14C**). Importantly, RNA-seq data revealed that *Nat10* knockout is associated with down-regulation of *Pgr* mRNA (**Fig. S15A-C and Table S4**). The loss of NAT10 at the protein level was confirmed via western blot analysis (**Fig. 5C**). Immunohistochemical analysis of NAT10 deletion efficiency demonstrated efficient deletion in stromal cells but preservation in luminal and glandular epithelial cells (**Fig. 5D**). Under normal circumstances, NAT10 deletion in epithelial cells should be complete before PND14 [23]. To elucidate why NAT10 remains intact in epithelial cells but is deleted in stromal cells, both cell types were isolated from *Nat10*^f/f^ mice (with wild-type mice as controls) and infected with an adenovirus expressing Cre (Adv-Cre) to inactivate the floxed alleles. *Nat10* knockout was evident at the protein level 4 days post-infection. Our findings revealed that *Nat10* knockout had no impact on cell apoptosis in either epithelial or stromal cells. Notably, *Nat10*-knockout epithelial cells failed to proliferate, whereas *Nat10*-knockout stromal cells exhibited only reduced proliferation (**Fig. S16A-B**). These data suggest that the persistence of NAT10 expression in uterine epithelium of *Nat10*^d/d^ mice is probably a result of the selective growth of NAT10-positive epithelial cells.

**Figure 5.**
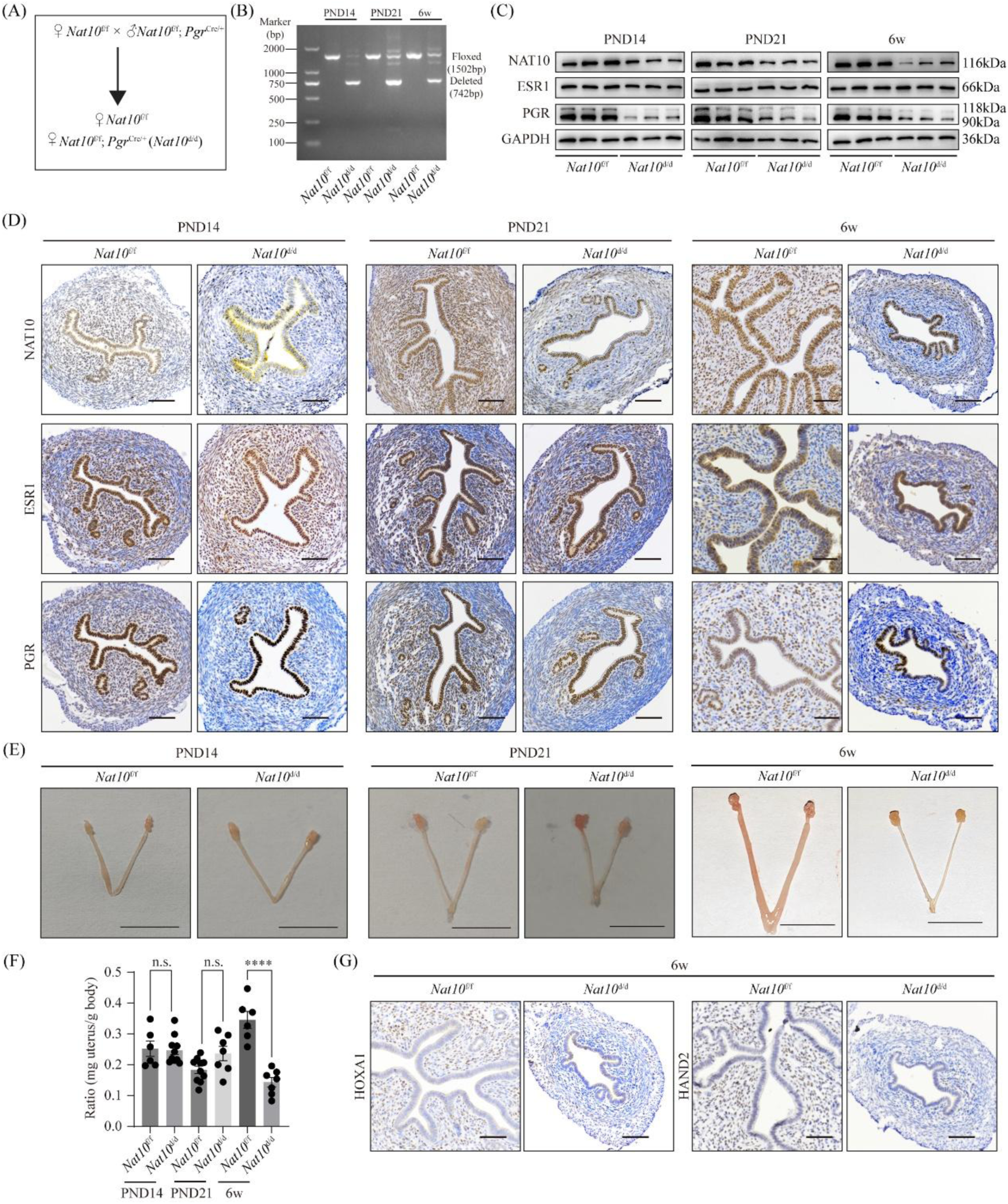
Pgr-Cre-mediated deletion of *Nat10* in mice results in impaired pubertal uterine development. (A) Diagram of breeding scheme. Uterine-specific *Nat10* deletion (*Nat10*^d/d^) mice were generated by crossing *Nat10*^f/f^ mice with a *Pgr*-Cre driver (*Pgr*^Cre/+^). (B) PCR analysis showing the knockout efficiency of *Nat10*^d/d^ mice at DNA level in the uterus on postnatal days 14 (PND14), 21 (PND21), and at 6 weeks (6w). (C) Western blot analysis of NAT10, ESR1 and PGR protein levels in the uterus of *Nat10*^d/d^ mice on PND14, PND21 and 6w. (D) Immunohistochemistry analysis of NAT10, ESR1 and PGR protein levels in the uterus of *Nat10*^d/d^ mice on PND14, PND21 and 6w. Bar=50 μm. (E) Representative photos of the uterus from *Nat10*^d/d^ mice on PND14, PND21 and 6w. (F) Statistical analysis of uterine weight in *Nat10*^d/d^ mice on PND14, PND16 and 6w. Data are presented as mean ± SEM. ****, P < 0.0001. (G) Immunohistochemistry staining of HOXA11 and HAND2 in the uterus *Nat10*^d/d^ mice at 6w. Bar=50 μm.

Gross morphological analysis of the uteri of *Nat10*^d/d^ mice revealed that, prior to puberty at PND14 and PND21, there were no alterations in their overall size or weight. However, a significant reduction in these parameters was evident after puberty at 6w (**Fig. 5E-F**). To further investigate, we assessed a range of protein markers, including the pan epithelial cell marker KRT8, luminal epithelial cell marker TACSTD2, glandular epithelial cell marker FOXA2, stromal cell marker VIM, smooth muscle cell marker ACTA2, endothelial cell marker PECAM1, and macrophage marker ADGRE1. This analysis demonstrated a reduced stroma layer and a decrease in the number of uterine glands in the *Nat10*^d/d^ uterus at 6w (**Fig. S17**). To examine changes in proliferation and apoptosis, immunostaining was performed for the proliferative marker MKI67 and the apoptotic enzyme cleaved CASP3. Our results indicated a scarcity of MKI67-positive cells in the stromal layer of *Nat10*^d/d^ uteri, in contrast to the numerous MKI67-positive cells observed in the stromal layer of *Nat10*^f/f^ uteri at 6w. Notably, no cleaved CASP3-positive cells were detected in either *Nat10*^f/f^ or *Nat10*^d/d^ uteri at all timepoints examined (**Fig. S17**). Additionally, we found that while ESR1 protein levels remained unchanged, PGR protein levels were significantly downregulated in the stromal cells from *Nat10*^d/d^ uteri at PND14, PND21 and 6w (**Fig. 5D**). Consistent with this finding, PGR target genes HAND2 and HOXA11 were also downregulated at 6w (**Fig. 5G**). These results suggest that the deletion of *Nat10* in stromal cells leads to decreased PGR expression and reduced cellular proliferation.

### Loss of uterine stromal NAT10 leads to infertility due to impaired uterine receptivity and failed decidual response

To investigate the significance of NAT10 in female fertility among adults, we conducted a 6-month fertility test by pairing *Nat10*^d/d^ and *Nat10*^f/f^ female mice with fertile wild-type male mice of the same genetic strain. Our findings revealed that *Nat10*^d/d^ female mice were infertile, whereas *Nat10*^f/f^ control female mice maintained normal fertility (**Fig. 6A**). These observations underscore the indispensable role of NAT10 in achieving successful pregnancy.

**Figure 6.**
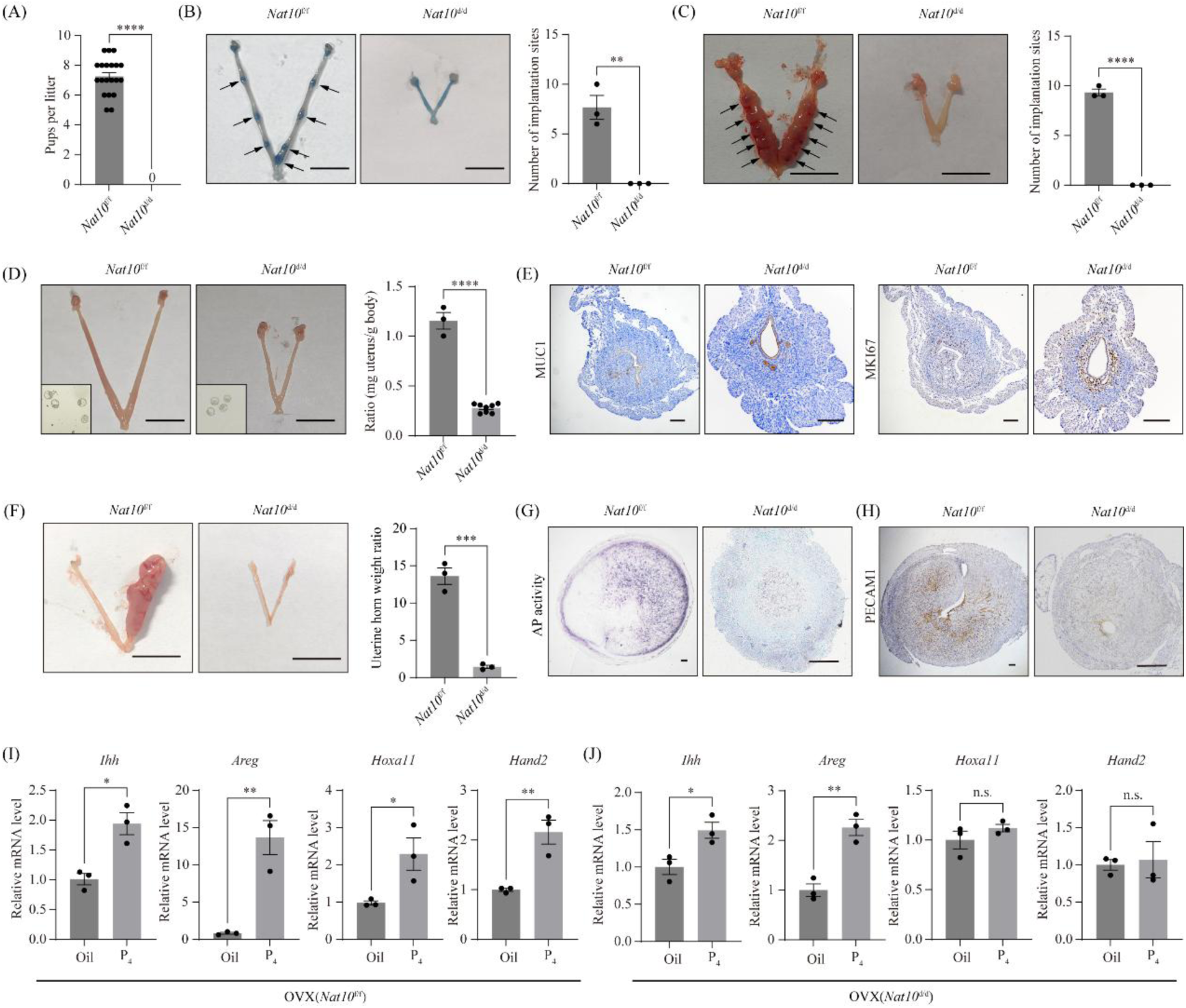
Deletion of *Nat10* in mouse uterus leads to infertility in adults. (A) Comparison of litter sizes for 3 *Nat10*^d/d^ mice and 3 *Nat10*^f/f^ mice over a 6-month period. Data are presented as mean ± SEM. ****, P<0.0001. (B) Bar plot showing the number of embryo implantation sites in *Nat10*^d/d^ and *Nat10*^f/f^ mice on gestational day 5 (GD5). Implantation sites were visualized by the blue dye injection method. Arrowheads mark implantation sites. Bar = 1 cm. Data are presented as mean ± SEM. **, P<0.01. (C) Bar plot showing the number of embryo implantation sites in *Nat10*^d/d^ and *Nat10*^f/f^ mice on GD8. Bar = 1 cm. Data are presented as mean ± SEM. ****, P<0.0001. (D) Bar plot showing the uterine weight in *Nat10*^d/d^ and *Nat10*^f/f^ mice on GD4. Bar = 1 cm. Data are presented as mean ± SEM. ****, P<0.0001. (E) Immunohistochemistry analysis of MUC1 and MKI67 protein levels in the uterus of *Nat10*^d/d^ and *Nat10*^f/f^ mice on GD4. Bar=100 μm. (F) Bar plot showing the weight ratio between stimulated uterine horn to the un-stimulated uterine horn in *Nat10*^d/d^ mice and *Nat10*^f/f^ mice following artificial decidualization. Bar = 1 cm. Data are presented as mean ± SEM. ***, P<0.001. (G) Alkaline phosphatase staining of the stimulated uterine horn from *Nat10*^d/d^ and *Nat10*^f/f^ mice. Bar=100 μm. (H) Immunohistochemistry staining of PECAM1 in the stimulated uterine horn from *Nat10*^d/d^ and *Nat10*^f/f^ mice. Bar=100 μm. (I-J) Quantitative RT-PCR analysis of P4 target genes in the uterus of ovariectomized mice for *Nat10*^f/f^ (I) and *Nat10*^d/d^ (J). Data are presented as means ± SEM. *, P < 0.05; **, P < 0.01.

To identify the cause of infertility in *Nat10*^d/d^ mice, we examined the pregnancy status of *Nat10*^d/d^ mice at different timepoints. Notably, on GD5 and GD8, *Nat10*^d/d^ mice exhibited a complete absence of implantation sites (**Fig. 6B-C**). However, we were able to recover embryos from the uterus of *Nat10*^d/d^ mice on GD4 (**Fig. 6D**). We next investigated whether the infertility of *Nat10*^d/d^ females was due to an ovarian defect. Through immunohistochemical analysis, we observed that the expression of NAT10 remained unaffected in the ovaries of *Nat10*^d/d^ mice on GD4 (**Fig. S18A**). Furthermore, the critical steroid biosynthetic enzymes, CYP11A1 (P450scc) and HSD3B2, were normally expressed in the corpus luteum of these mice (**Fig. S18B**). Additionally, serum levels of E_2_ and P_4_ were comparable between *Nat10*^d/d^ and control *Nat10*^f/f^ female mice on GD4 (**Fig. S18C**), further supporting normal ovarian activity. We then examined possible defects in uterine function. On GD4, the uterine size and weight of *Nat10*^d/d^ mice were significantly reduced compared to those of *Nat10*^f/f^ mice (**Fig. 6D**). Histological analysis demonstrated a reduced stroma layer and a decrease in the number of endometrial glands in the *Nat10*^d/d^ uterus (**Fig. S19**), which can be traced back to pubertal uterine development (**Fig. S17**). Next, we evaluated uterine receptivity markers. Specifically, the loss of the anti-adhesive glycoprotein mucin 1 (MUC1) on the entire luminal epithelium serves as an indicator of uterine receptivity in mice. Our immunohistochemical analysis showed abundant MUC1 protein on the luminal epithelium of *Nat10*^d/d^ mice, whereas it was absent on the luminal epithelium of *Nat10*^f/f^ mice. The cessation of luminal epithelium proliferation is pivotal for establishing uterine receptivity. Immunohistochemical staining for MKI67, a marker of proliferative cells, indicated persistent proliferation in the luminal epithelium of *Nat10*^d/d^ mice on GD4 (**Fig. 6E**). It is noteworthy that the deletion of NAT10 in uterine stromal cells is linked to decreased expression of PGR (**Fig. S20A**), mirroring the changes observed during pubertal uterus development. Since uterine receptivity is strictly regulated by P_4_, we examined the impact of NAT10 loss on the expression of P_4_ target genes in the uteri of mice at GD4. Quantitative RT-PCR analysis showed significant downregulation of stromal target genes *Hoxa10* and *Hand2* in *Nat10*^d/d^ uteri compared to *Nat10*^f/f^ uteri, while epithelial target genes *Ihh* and *Areg* remained unchanged (**Fig. S20B**). The downregulation of HOXA11 and HAND2 was further validated through immunohistochemical analysis (**Fig. S20C**). These findings suggest that the deletion of NAT10 in uterine stromal cells disrupts uterine receptivity by impeding the P_4_ response.

Based on our hESC studies, we hypothesized that NAT10-deficiency would incur decidualization defects at pregnancy in mice. To investigate whether *Nat10* deletion had an impact on decidualization, we employed an artificial decidualization model. Notably, the uterus of *Nat10*^f/f^ mice displayed a robust decidual response, whereas absence of such a response was observed in *Nat10*^d/d^ mice (**Fig. 6F**). Alkaline phosphatase activity, a well-recognized indicator of stromal cell differentiation, was evident in the uterus of *Nat10*^f/f^ mice but not in that of *Nat10*^d/d^ mice following artificial decidualization (**Fig. 6G**). During decidualization, stromal cells secrete various factors that promote vascular development. Consequently, we investigated the extent of angiogenesis by immunostaining for platelet/endothelial cell adhesion molecule 1 (PECAM1), a marker of endothelial cells. As anticipated, PECAM1 immunostaining was significantly reduced in the uterus of *Nat10*^d/d^ mice after artificial decidualization (**Fig. 6H**). Moreover, the compromised decidualization in *Nat10*^d/d^ mice was further corroborated by quantitative RT-PCR analysis of decidualization marker genes, including bone morphogenetic protein 2 (*Bmp2*), Wnt family member 4 (*Wnt4*), and prolactin family 8 subfamily A member 2 (*Prl8a2*) (**Fig. S21A**). Notably, deletion of NAT10 in uterine stromal cells is associated with reduced PGR expression, along with the down-regulation of its targets HOXA11 and HAND2 (**Fig. S21B-C**). In summary, these findings indicate that the deletion of *Nat10* leads to compromised decidualization.

To further establish that the decreased PGR expression in uterine stromal cells of *Nat10*^d/d^ mice attenuates P_4_ signaling, we conducted an assay to measure the expression of P_4_ target genes in the uterus of ovariectomized *Nat10*^d/d^ mice and *Nat10*^f/f^ mice, 12 hours after they were administered either P_4_ (1 mg) or a vehicle control (oil). Our findings revealed that the *Nat10*^f/f^ mice exhibited an elevation in the expression of P_4_ target genes, specifically *Ihh*, *Areg*, *Hoxa11*, and *Hand2* (**Fig. 6I**). Conversely, the *Nat10*^d/d^ mice demonstrated a complete absence of P_4_-induced upregulation of stromal-specific genes *Hoxa11* and *Hand2*. It is worth noting that the expression of epithelial genes *Ihh* and *Areg* was either unaffected or only partially affected in the *Nat10*^d/d^ mice (**Fig. 6J**). These observations suggest that P_4_ signaling is specifically compromised within the stromal compartment of the *Nat10*^d/d^ uterus.

### Pharmacological inhibition of NAT10 activity in wild-type mice leads to impaired implantation and decidualization

Given the abnormal uterine structure observed in adult *Nat10*^d/d^ mice, it is conceivable that the infertility phenotype may not solely arise from the absence of NAT10 during the implantation phase, but rather due to uterine structural abnormalities that develop during pubertal growth. To address this issue, we created *Nat10*^d/+^ mice (*Pgr*^Cre/+^; *Nat10*^f/+^), in which NAT10 is partially knocked out in the uterus (**Fig. S22A**). A notable reduction in NAT10 expression was observed in the uterus at both the mRNA and protein levels (**Fig. S22B-D**). Importantly, this downregulation of NAT10 was accompanied by a concurrent decrease in PGR expression (**Fig. S22C and E**). In contrast to *Nat10*^d/d^ mice, *Nat10*^d/+^ mice displayed normal uterine development, characterized by typical stromal thickness and a normal number of glands (**Fig. S22F**). Upon close inspection, Nat10^d/+^ females are normal in uterine receptivity (**Fig. S22G**), implantation rate (**Fig. S22H**) and decidualization response (**Fig. S22I**). Moreover, the fertility test revealed that *Nat10*^d/+^ females possessed normal fertility (**Fig. S22J**).

A plausible explanation for the normal fertility observed in mice with partial *Nat10* knockout is that the reduced gene dose in these animals is insufficient to elicit a notable phenotypic effect. Remodelin is a specific inhibitor of NAT10 [24]. In hESCs, we observed that cell morphology appeared normal following treatment with up to 50 µM of Remodelin (**Fig. S23A**). Notably, a concentration of 30 µM of Remodelin was sufficient to effectively inhibit PGR expression (**Fig. S23B**) and suppressing in vitro decidualization (**Fig. S23C-D**). These results provided us with encouraging preliminary evidence to proceed with in vivo studies. WT mice were orally administered Remodelin at 100 mg/kg daily from GD3 to GD4, a dosage widely recognized as safe and effective for NAT10 inhibition [25, 26]. Implantation was examined in uteri on GD5 (**Fig. 7A**). Compared to the control group, the number of implantation sites was significantly reduced in the Remodelin-treated animals **(Fig. 7B**). We conducted serial sectioning and PTGS2 staining to confirm each suspicious implantation site with a blurred appearance (**Fig. 7C**). Moreover, we found that PGR expression was downregulated at the implantation sites in Remodelin-treated mice compared to control mice (**Fig. 7D**). These results suggest that inhibiting NAT10 impairs the implantation process.

**Figure 7.**
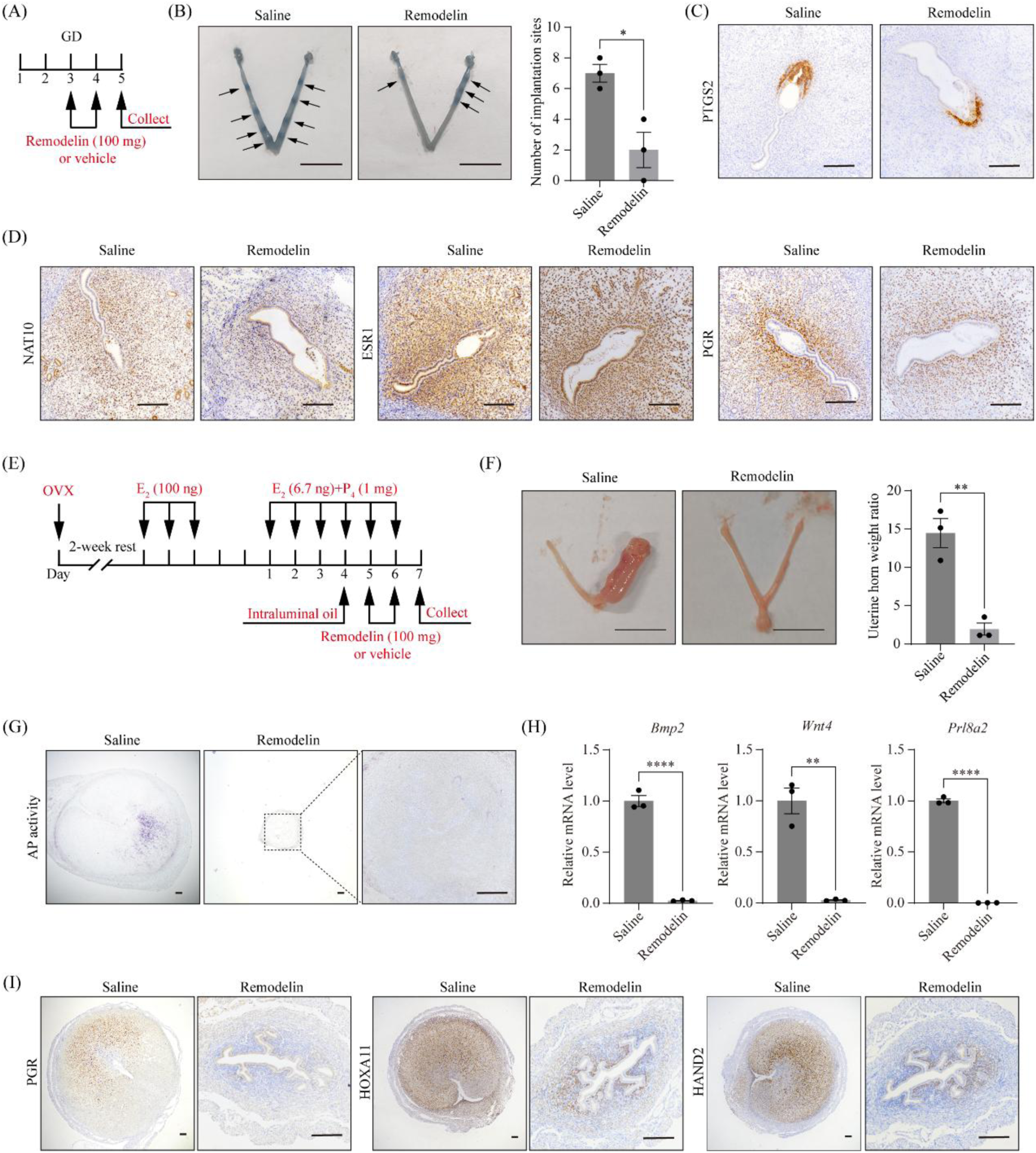
Remodelin administration impairs implantation and decidualization in vivo. (A) The experimental protocol for Remodelin administration in pregnant mice. GD, gestational day. (B) The impact of Remodelin on implantation in mice. Implantation sites were visualized by the blue dye injection method on GD5. Implantation sites are marked by arrowheads. Data are presented as means ± SEM. *, P < 0.05. (C-D) Immunohistochemistry staining of PTGS2, NAT10, ESR1 and PGR at the implantation sites of the uterus from Remodelin-treated and control mice. Bar=100 μm. (E) The experimental protocol for Remodelin administration in ovariectomized mice undergoing artificial decidualization. (F) Bar plot comparing the weight ratio between stimulated horn to the un-stimulated horn for Remodelin-treated and control mice. Data are presented as means ± SEM. **, P < 0.01. (G) Representative images showing alkaline phosphatase staining of the stimulated uterine horn in Remodelin-treated and control mice. Bar=100 μm. (H) Bar plot depicting the quantitative RT-PCR analysis of decidualization marker genes in the stimulated uterine horn of Remodelin-treated and control mice. Data are presented as means ± SEM. **, P < 0.01; ****, P < 0.0001. (I) Representative images of immunohistochemical staining for PGR, HOXA11, and HAND2 in the stimulated uterine horn of Remodelin-treated and control mice. Bar=100 μm.

Since implantation is regulated by three factors: the ovary, embryo, and uterus, we conducted further experiments to exclude the potential impact of Remodelin treatment on the ovary and embryo (**Fig. 7E**). To exclude the ovary factor, adult female mice were ovariectomized and rested for 2 weeks to clear endogenous ovarian hormones. Following this, the mice were primed with steroid hormones to induce an artificial receptive stage. To exclude the embryo factor, sesame oil was injected into the uterine lumen to mimic the presence of an embryo. Our findings revealed that, in control mice, the decidual response was evident in a manner similar to that observed during natural embryo implantation. However, in mice treated with Remodelin, this response was impaired, as indicated by the size and weight of the uterine horn (**Fig. 7F**). Impaired decidualization was confirmed by the absence of alkaline phosphatase activity (**Fig. 7G**) and reduced expression of decidualization marker genes *Prl8a2*, *Bmp2*, and *Wnt4* (**Fig. 7H**). As expected, inhibition of NAT10 by Remodelin was associated with decreased PGR expression and downregulation of its targets HOXA11 and HAND2 (**Fig. 7I**). These findings suggest that inhibition of the NAT10-PGR axis in the mouse uterus significantly impairs decidualization.

## Discussion

Studies have established that ac^4^C modification is crucial in numerous biological processes [8]. In this study, we revealed that knockdown of NAT10, the enzyme responsible for ac^4^C modification, in human endometrial stromal cells (hESCs) impeded the process of decidualization. Notably, this impairment was primarily attributed to the reduced expression of PGR. Mechanistically, we discovered that *PGR* mRNA is a direct substrate for NAT10-mediated ac^4^C modification, and that this modification within the coding sequence (CDS) of *PGR* mRNA enhances its stability. In mice, conditional knockout of *Nat10* using *Pgr*-Cre resulted in defective pubertal uterine development and infertility. In contrast, heterozygous conditional knockout mice exhibited decreased NAT10 expression but did not display uterine defects. Additionally, administration of Remodelin, a NAT10 inhibitor, maintained uterine structures but disrupted the P_4_/PGR signaling pathway in adult mice, resulting in impaired uterine function during pregnancy. This study elucidates the physiological role of ac^4^C modification in the uterus during pregnancy.

In our study, we observed that NAT10 undergoes downregulation during the process of decidualization in hESCs in vitro. This finding is in accordance with our observation that NAT10 is downregulated in the human endometrium during the late secretory phase compared to the middle secretory phase of the menstrual cycle. Similarly, NAT10 downregulation was also noted during decidualization in mouse endometrial stromal cells (mESCs) in vitro and in the basal decidua of the mouse uterus from gestation day 8 (GD8) to GD18 in vivo. We further demonstrated that NAT10 downregulation is mediated by both MPA and cAMP. In hESCs, knockdown of NAT10 at any stage of decidualization adversely affected the outcome of the process, and overexpression of NAT10 similarly impaired decidualization. As a target of NAT10, PGR also undergoes downregulation during in vitro decidualization in hESCs, aligning with previous research [27]. PGR downregulation was also observed in the human endometrium during the secretory phase [22]. Our results indicate that PGR is downregulated by MPA but upregulated by cAMP, corroborating findings from a previous study [28]. Both knockdown and overexpression of PGR impairs decidualization in hESCs. In mice, both PGR knockout and overexpression resulted in failures in uterine receptivity and decidualization [29, 30], suggesting that PGR has a dual role in decidualization, being both beneficial and harmful. A plausible explanation for this dual effect is that PGR itself inhibits decidualization [27] while its downstream targets may promote decidualization [31, 32]. Current research on decidualization has predominantly focused on the role of upregulated genes [33]. However, our study demonstrates that programmed downregulation of NAT10 and PGR is crucial for this process. These findings may provide valuable insights into the mechanisms underlying decidualization, expanding our understanding of the intricate interplay between gene expression pattern and decidualization.

NAT10 features both an acetyltransferase domain and an RNA-binding domain. It has been shown to exhibit catalytic activity towards both RNA and protein [19]. Our study, employing dot blot analysis, revealed that RNA is the predominant substrate for NAT10 compared to proteins in hESCs. NAT10 has the capability to install ac^4^C modifications on various RNA types, including tRNA, rRNA, and mRNA [12]. Specifically, the acetylation of rRNA by NAT10 necessitates collaboration with small nucleolar RNA (snoRNA) U13, whereas acetylation of tRNA requires assistance from a specific RNA adapter protein known as THUMPD1. In rRNA, the presence of ac^4^C is pivotal for the formation of the ribosome 18S subunit. The ac^4^C modification in tRNA enhances its stability, translation efficiency, and fidelity. Given the broad impact of ac4C modifications on tRNA and rRNA on virtually all genes, we have a particular focus on ac^4^C modifications in mRNA, which may function in a gene-specific manner. Through the application of acRIP-seq, we discovered that PGR mRNA undergoes ac^4^C modification in both human and mouse endometrium. Using bioinformatic tools and a reporter system, we identified a conserved ac^4^C site within the CDS of *PGR* mRNA across humans and mice. It is established that ac^4^C modifications in the CDS enhance mRNA stability and translation efficiency [10]. Employing an mRNA stability assay, we validated that ac^4^C modification indeed impacts *PGR* mRNA stability. Notably, in our study of *PGR* mRNA, we found that NAT10 knockdown-induced downregulation of PGR is more pronounced at the mRNA level than at the protein level, leading us to conclude that ac4C modification primarily affects mRNA stability rather than translation efficiency. In contrast to m^6^A, which functions through specific reader proteins, the function of ac4C appears to be driven exclusively by its structural influence [34]. In ac^4^C modification, an acetyl group (-COCH_3_) is covalently linked to the *N*^4^ nitrogen atom via an amide bond, reducing cytosine polarity and enhancing its hydrophobicity, thus enhancing overall stability. Meanwhile, ac^4^C disrupts standard C-G base pairing, as the unmodified *N*^4^ amino group engages in typical hydrogen bonding. When ac^4^C is positioned at wobble sites within the mRNA CDS, it is believed to enhance translation efficiency by modifying the interaction between mRNA and tRNAs. However, in our study, ac^4^C sites in the *PGR* CDS were not located at wobble positions, which may explain why translation efficiency was not enhanced in our case. Collectively, this study has uncovered that NAT10 stabilizes *PGR* mRNA by installing ac^4^C modifications within its CDS region. Together with our previous research demonstrating that METTL3-mediated m^6^A modification serves as a crucial regulator of *PGR* translation in the uterus [17], our study underscores the significance of RNA modifications in regulating PGR expression, uterine function, and fertility.

In this study, we employed three mouse models to delve into the role of NAT10 in the uterus during pregnancy in vivo. In the *Nat10* conditional knockout (cKO) model, we observed that the uterine deletion of *Nat10* occurred only in stromal cells but not in epithelial cells. Using adenovirus expressing Cre recombinase (Adv-Cre), we further discovered that proliferation ceased in epithelial cells upon *Nat10* knockout, while proliferation was merely slowed in stromal cells. Considering that Cre activity may not be 100% efficient, the preserved expression of NAT10 in the uterine epithelium of cKO mice is likely due to the selective growth of NAT10-positive epithelial cells. This interpretation is supported by a previous study showing initial Ctcf ablation in uterine epithelium via *Pgr*-Cre with re-emergence during puberty [35]. Notably, previous studies have reported that NAT10 regulates centrosome physiological activities through its protein acetylation activity at various mitotic stages, thereby playing a role in cell division [36, 37]. The *Nat10* cKO resulted in impaired pubertal uterine development, characterized by a reduced stroma layer and a decrease in the number of endometrial glands. It is important to note that thin stroma and reduced gland number are likely phenotypes independent of PGR, as mice with *Pgr* knockout or *Pgr-A* knockout exhibit normal uterine structure [29, 38]. Thin stroma has been implicated in infertility in humans [39], and the absence of glands also causes infertility in mice [40]. Therefore, although infertility is observed in *Nat10* cKO mice, we cannot definitively attribute this infertility solely to NAT10. There is a possibility that the infertility arises from uterine structural abnormalities rather than a direct involvement of NAT10. To address this issue, we generated heterozygous *Nat10* cKO mice, which turned out to exhibit normal uterine structure and fertility. In these mice, both NAT10 and PGR are downregulated. However, the expression levels of PGR target genes, such as HOXA11 and HAND2, remain unchanged. This observation suggests that the partial *Nat10* cKO in these mice may not be sufficient to elicit a significant phenotypic effect. To bridge the gap between heterozygous and homozygous cKO conditions, we employed Remodelin, an inhibitor of NAT10, in adult female mice, to mimic an intermediate gene dosage. The mice were treated with Remodelin for two consecutive days, and the impacts on implantation and decidualization were assessed on the third day. We assumed that the uterine structure would remain undisturbed due to two reasons: (1) the brief duration of Remodelin exposure and (2) the nature of NAT10 inhibition by Remodelin, which, unlike genetic knockout, may have a more limited impact on cell proliferation, akin to siRNA knockdown. Through this approach, we demonstrated that the NAT10-PGR axis plays a crucial role in uterine function during implantation and decidualization. Collectively, this study demonstrated that the antagonism of NAT10 affects uterine phenotypes in a gene-dose-dependent manner. Our findings hold significant clinical importance, as we observed a downregulation of NAT10, rather than a complete knockout, in the stromal compartment of the endometrium in patients with recurrent pregnancy loss (RPL). Based on our mouse model data, this downregulation of NAT10 may have minimal impact on cell proliferation but could lead to decreased PGR expression and impaired decidualization. Further investigation is required to confirm these findings. Our study offers potential therapeutic insights for the treatment of RPL.

In summary, this study contributes to a more comprehensive understanding of P_4_ signaling through the identification of a role of NAT10 and ac^4^C modifications in orchestrating PGR expression and thereby ensuring optimal uterine function and fertility.

## Materials and Methods

### Human endometrial sample collection

The patients enrolled in this study were recruited from Reproductive Medicine Center of the First Affiliated Hospital of Sun Yat-sen University in China. Informed consent was obtained from all participants and this study was approved by the Ethical Committee of the First Affiliated Hospital of Sun Yat-sen University (No. 2018-266). Normal fertile participants who had no apparent endometrial pathology and had a confirmed clinical pregnancy after ET were selected. Detailed information of patients is described in our previous study [17].

### Isolation and culture of human endometrial stromal cells (hESCs)

Three healthy fertile participants, who exhibited no signs of endometrial pathology and had a clinically confirmed pregnancy following embryo transfer, were selected for this study. Prior to commencing the study, it was approved by the Ethical Committee of the First Affiliated Hospital of Sun Yat-sen University (No. 2018-266). All participants provided their informed consent. The detailed characteristics of these patients are summarized in **Table S5**. The endometrial tissues were dissected into pieces and then underwent digestion using type I collagenase (Gibco) for 1 h. Subsequently, the endometrial epithelial cells and stromal cells were separated using membrane filters (100 µm and 40 µm cell filters, Corning). The hESCs were cultured in DMEM/F12 medium (Gibco), supplemented with 10% charcoal-stripped fetal bovine serum (cFBS, VivaCell). To induce decidualization, the cells were treated with 0.5 mM 8-Br-cAMP (Sigma) and 1 μM medroxyprogesterone acetate in 2% cFBS.

### Mice

The *Nat10*^f/f^ mice (Cat. No. NM-CKO-2117429) and *Pgr*^Cre/+^ mice (Cat. No. NM-KI-200117) were acquired from Shanghai Model Organisms Center, Inc., China. The *Nat10*^d/d^ and *Nat10*^d/+^ mice were generated through the crossing of *Nat10*^f/f^ mice with *Pgr*^Cre/+^ mice. The *Nat10*^f/f^ littermates served as the wild-type control group. Adult female mice were bred with either fertile males or vasectomized males to induce pregnancy or pseudo-pregnancy, respectively. The day of vaginal plug detection was designated as gestational day 1 (GD1). For the artificial decidualization experiments, a 20 μl volume of sesame oil (Sigma) was injected into one uterine horn of the pseudo-pregnant mice on GD4, and uterine samples were collected on GD8. All animal procedures conducted were approved by the Institutional Animal Care and Use Committee of South China Agricultural University, under approval number 2018B028.

To examine the effect of Remodelin on embryo implantation in vivo, wild-type pregnant mice were orally administered Remodelin (100 mg/kg body weight; V2042, InvivoChem, Guangzhou, China) on GD3-4. Implantation sites were visualized by the blue dye method on GD5. To examine the effect of Remodelin on decidualization in vivo, an artificial decidualization model with ovariectomized mice was employed. Adult female mice were ovariectomized. After a 2-wk rest, mice were subcutaneously injected with 100 ng E_2_ (Sigma) for 3 days. After 2-day rest, 6.7 ng E_2_ (Sigma) plus 1 mg P_4_ (Sigma) was injected daily. On day 4 of E_2_ and P_4_ injection, 10 μL of sesame oil (Sigma) was injected into the lumen of one uterine horn to induce decidualization. The contralateral horn serving as control. Mice were orally administered Remodelin (100 mg/kg body weight) on days 5-6. Uterine samples were collected on day 7.

### Primary culture of mouse endometrial stromal cells (mESCs)

The mESCs were isolated and cultured with slight modifications to a previously described method [41]. Briefly, uterine horns from adult mice on GD4 were dissected into small fragments. These fragments were then submerged in Hank’s balanced salt solution (HBSS) containing 6 mg/ml dispase and 25 mg/ml trypsin for 1h at 4°C. Following this, the tissues were incubated for an additional 1h at room temperature and then for 10 min at 37°C. After removing the endometrial epithelial clumps, the remaining tissues were incubated once more in HBSS containing 0.5 mg/ml collagenase at 37°C for 30 min. To obtain stromal cells, the digested cells were filtered through a 70-μm mesh. The isolated cells were seeded onto 60-mm dishes at a density of 5 × 10^5^ cells per dish and cultured in a mixture of phenol red-free Dulbecco’s modified Eagle’s medium and Ham F-12 nutrient medium (1:1 ratio; DMEM/F12; Gibco), supplemented with 10% charcoal-stripped fetal bovine serum (cFBS; Biological Industries) and antibiotics. After an initial 2h incubation, the medium was replaced with a phenol red-free DMEM/F-12 containing 1% cFBS, 10 nM estradiol (E_2_), and 1 μM progesterone (P_4_) to induce in vitro decidualization.

### Quantitative RT-PCR

Total RNA was extracted using the TRIzol reagent (Invitrogen). Subsequently, 1 μg of this purified RNA was utilized to synthesize cDNA with the HiScript III RT Super Mix (Vazyme). Quantitative PCR was conducted using the ChamQ SYBR qPCR Master Mix (Vazyme) on the Applied Biosystems 7300 Plus (Life Technologies). The expression data were normalized against the expression of *Rpl7* as a reference gene. The comprehensive list of primer sequences employed in this study is provided in **Table S6**.

### Western blot and protein dot blot

Uterine tissues were homogenized, and cultured cells were lysed using Radio-Immunoprecipitation Assay (RIPA) buffer supplemented with protease inhibitors (Roche). For western blot, the total protein extracts were then resolved on a 10% SDS-PAGE gel and transferred onto a PVDF membrane (Millipore). For protein dot blot analysis, the protein lysates were loaded onto a PVDF membrane and fixed to the membrane by drying at 37°C for 30 min. Prior to antibody probing, the membrane was blocked with 5% skim milk in Tris-buffered saline with Tween 20 (TBST). Subsequently, the membrane was incubated overnight at 4°C with the primary antibody. After thorough washing, the membrane was incubated with a horseradish peroxidase (HRP)-conjugated secondary antibody for 1h at room temperature. The immunoreactive signals were visualized using the ECL chemiluminescent kit (Amersham Biosciences). GAPDH bands were utilized as the loading control. Band intensities were quantitatively analyzed using the ImageJ software version 1.54i [42]. The primary antibodies employed in this study are listed in **Table S7**.

### RNA dot blot

The mRNA was harvested from total RNA using the Dynabeads® mRNA DirectTM Purification Kit (Invitrogen). After denaturing the RNA samples at 95 °C, they were immediately chilled on ice. Subsequently, the RNA samples were spotted onto an Amersham^TM^ Hybond^TM^-N+ membrane (GE Healthcare) and affixed to the membrane through UV cross-linking. To reduce non-specific binding, the membrane was blocked with 5% non-fat milk for 1 h. Following this, the membrane was incubated overnight at 4°C with an m6A-specific antibody. After thorough washing with PBST, the membrane was incubated with the secondary antibody, HRP-conjugated goat anti-rabbit IgG, for 1 h at room temperature. The immunoreactive signals were visualized using an ECL chemiluminescent kit (Amersham Biosciences). Methylene blue staining was performed as a loading control. Dot intensities were quantitatively analyzed using the ImageJ software v1.54i [42].

### Immunofluorescence

Cellular fixation was performed using 4% paraformaldehyde (P6148, Sigma) for 15 minutes, followed by permeabilization with 0.1% Triton X-100 (T8787, Sigma) for 10 minutes. Non-specific binding was blocked with 10% goat serum incubation at 37°C for 1 hour. Primary antibody targeting the protein of interest was applied overnight at 4°C. On the following day, samples were incubated with species-specific secondary antibodies (Jackson ImmunoResearch) for 40 minutes at room temperature. Immunofluorescence staining with phalloidin was performed according to the instructions provided in the kit from Beyotime (C2201S). A complete list of primary antibodies with corresponding dilutions and suppliers is provided in **Table S7**.

### RNA-seq

Uterine segments were frozen in liquid nitrogen and stored at -80 °C until use. Total RNA extraction, cDNA library construction and high-throughput DNA sequencing were executed following our previously established protocols. After quality control, the sequence data were aligned to the mouse genome (UCSC mm10) using HISAT v2.2.1 [43]. Gene-level expression values were derived from the aligned sequences with Cufflinks v2.2.1 [44]. Differentially expressed genes were selected based on the following criteria: fold change > 2 and adjusted P-value < 0.05. To gain insights into the functional roles of differentially expressed genes, gene ontology analysis was performed using the DAVID online tools with knowledgebase v2024q4 [45]. Redundant enriched terms were manually eliminated.

### acRIP-seq

Purified mRNA was fragmented in fragmentation buffer (Invitrogen, AM8740) by heating it at 94℃ for 5 min. The ac^4^C antibody immunoprecipitated RNA fragments, along with the input control, were prepared using the GenSeq® ac^4^C RIP kit (GS-ET-005, GenSeq Inc., China). The acRIP-Seq libraries were generated with the Next® Ultra™ II Directional RNA Library Prep Kit (NEB) and sequenced on the Illumina HiSeq 2500 system. After quality control, the sequence data were aligned to the mouse genome (UCSC mm10) with HISAT v2.2.1 [43]. For visual inspection, the sequence alignment on the genome was displayed using the IGV tool v2.14.0 [46]. Peak calling was performed with MACS software v3.0.0a7 with q-value < 0.05 [47]. The metagene profile was analyzed using the R package Guitar v2.12.0 [48]. Motif analysis was carried out with HOMER v4.7 [49].

### RNA decay assay

Cells were plated in 12-well dishes and incubated overnight at 37 °C. For investigating NAT10-mediated mRNA decay, the cells were transfected with NAT10 siRNA or overexpression vector. After transfection for 48h, cells were treated with actinomycin D (A1410, Sigma) at a final concentration of 5 μg/ml to halt transcription. At various time intervals, total RNA was isolated from these cells and quantitative RT-PCR was performed to quantify the expression levels of *PGR* mRNA. The obtained data were normalized to the initial time point (t = 0) for comparison. The siRNAs and overexpression vectors used in this study are listed in **Table S8**.

### Immunohistochemistry

Paraformaldehyde-fixed paraffin-embedded uterine tissues were sliced into 5-μm sections. Heat-induced antigen retrieval was then conducted in a 10 mM citrate buffer (pH = 6.0) for a duration of 15 min. To eliminate endogenous peroxidase activity, the sections were treated with 3% H_2_O_2_ for 20 min. After blocking with 10% horse serum in PBS, sections were incubated with the primary antibody overnight at 4°C. After this, they were incubated with an HRP-conjugated secondary antibody for one hour at room temperature. The immunoreactive signals were visualized using DAB staining kit (Zhongshan Golden Bridge Biotechnology Co., Beijing). For contrast, the sections were counterstained with hematoxylin. Primary antibodies used in this study are listed in **Table S7**.

### Alkaline phosphatase staining

Frozen tissue sections, precisely sliced at 10 μm, were swiftly fixed in cold acetone for 15 minutes and subsequently washed thoroughly three times with PBS. To visualize alkaline phosphatase activity, the BCIP/NBT kit (Zhongshan Golden Bridge Biotechnology Co., Beijing) was employed, following the manufacturer’s recommended protocol. For counterstaining, a 1% solution of methyl green was utilized.

### Statistical analysis

Statistical analysis was performed using GraphPad Prism v10.0.0. The two-tailed unpaired Student’s t test was used for comparison of means of two groups. The one-way ANOVA with Tukey’s multiple comparison test was used for comparison of means of more than two groups. Data are presented as the mean ± SEM and P < 0.05 was considered statistically significant.

## Acknowledgements

This research was funded by National Natural Science Foundation of China (32370913 and 32070845), Guangdong Natural Science Funds for Distinguished Young Scholars (2021B1515020079), Innovation Team Project of Guangdong University (2019KCXTD001), Guangdong Special Support Program (2019BT02Y276), National Key R&D Program of China (2018YFA0801404), and Double first-class discipline promotion project (2023B10564003).

## Author contributions

Y.-W.X., S.-H.Y. and J.-L.L. designed research; S.T., D.-K.P., R.-F.J., Y.-Q.H., Q.-Y.Z., X.-Q.Y., A.-N.Z. and Y.-M.D. performed the experiments; S.T., Y.-W.X., S.-H.Y. and J.-L.L. analyzed the data. S.T., Y.-W.X., S.-H.Y. and J.-L.L. wrote the paper. All authors read and approved the final paper.

## Data availability statements

The sequencing data generated in this study are deposited in the Gene Expression Omnibus (GEO) under accession codes GSE293341, GSE293343, GSE293345, and GSE293346.

## References

1. Habiba M, Heyn R, Bianchi P, Brosens I, Benagiano G. The development of the human uterus: morphogenesis to menarche. Hum Reprod Update. 2021;27(1):1–26. Epub 2021/01/05. doi: 10.1093/humupd/dmaa036. PubMed PMID: 33395479.

2. Gellersen B, Brosens JJ. Cyclic decidualization of the human endometrium in reproductive health and failure. Endocr Rev. 2014;35(6):851–905. Epub 2014/08/21. doi: 10.1210/er.2014-1045. PubMed PMID: 25141152.

3. Muter J, Lynch VJ, McCoy RC, Brosens JJ. Human embryo implantation. Development. 2023;150(10). Epub 2023/05/31. doi: 10.1242/dev.201507. PubMed PMID: 37254877; PubMed Central PMCID: PMCPMC10281521.

4. Stocco C, Telleria C, Gibori G. The molecular control of corpus luteum formation, function, and regression. Endocr Rev. 2007;28(1):117–49. Epub 2006/11/02. doi: 10.1210/er.2006-0022. PubMed PMID: 17077191.

5. Cha J, Sun X, Dey SK. Mechanisms of implantation: strategies for successful pregnancy. Nat Med. 2012;18(12):1754–67. Epub 2012/12/12. doi: 10.1038/nm.3012. PubMed PMID: 23223073; PubMed Central PMCID: PMCPMC6322836.

6. Wu SP, Li R, DeMayo FJ. Progesterone Receptor Regulation of Uterine Adaptation for Pregnancy. Trends Endocrinol Metab. 2018;29(7):481–91. Epub 2018/05/01. doi: 10.1016/j.tem.2018.04.001. PubMed PMID: 29705365; PubMed Central PMCID: PMCPMC6004243.

7. Sun H, Li K, Liu C, Yi C. Regulation and functions of non-m(6)A mRNA modifications. Nat Rev Mol Cell Biol. 2023;24(10):714–31. Epub 2023/06/28. doi: 10.1038/s41580-023-00622-x. PubMed PMID: 37369853.

8. Wang Q, Yuan Y, Zhou Q, Jia Y, Liu J, Xiao G, et al. RNA N4-acetylcytidine modification and its role in health and diseases. MedComm (2020). 2025;6(1):e70015. Epub 2025/01/07. doi: 10.1002/mco2.70015. PubMed PMID: 39764566; PubMed Central PMCID: PMCPMC11702397.

9. Sas-Chen A, Thomas JM, Matzov D, Taoka M, Nance KD, Nir R, et al. Dynamic RNA acetylation revealed by quantitative cross-evolutionary mapping. Nature. 2020;583(7817):638-43. Epub 2020/06/20. doi: 10.1038/s41586-020-2418-2. PubMed PMID: 32555463; PubMed Central PMCID: PMCPMC8130014.

10. Arango D, Sturgill D, Alhusaini N, Dillman AA, Sweet TJ, Hanson G, et al. Acetylation of Cytidine in mRNA Promotes Translation Efficiency. Cell. 2018;175(7):1872–86 e24. Epub 2018/11/20. doi: 10.1016/j.cell.2018.10.030. PubMed PMID: 30449621; PubMed Central PMCID: PMCPMC6295233.

11. Ito S, Horikawa S, Suzuki T, Kawauchi H, Tanaka Y, Suzuki T, et al. Human NAT10 is an ATP-dependent RNA acetyltransferase responsible for N4-acetylcytidine formation in 18 S ribosomal RNA (rRNA). J Biol Chem. 2014;289(52):35724–30. Epub 2014/11/21. doi: 10.1074/jbc.C114.602698. PubMed PMID: 25411247; PubMed Central PMCID: PMCPMC4276842.

12. Wubulikasimu Z, Zhao H, Mao F, Zhao X. Emerging roles of RNA N4-acetylcytidine modification in reproductive health. Protein Cell. 2025. Epub 2025/02/19. doi: 10.1093/procel/pwaf013. PubMed PMID: 39969189.

13. Talbi S, Hamilton AE, Vo KC, Tulac S, Overgaard MT, Dosiou C, et al. Molecular phenotyping of human endometrium distinguishes menstrual cycle phases and underlying biological processes in normo-ovulatory women. Endocrinology. 2006;147(3):1097–121. Epub 2005/11/25. doi: 10.1210/en.2005-1076. PubMed PMID: 16306079.

14. Lai ZZ, Wang Y, Zhou WJ, Liang Z, Shi JW, Yang HL, et al. Single-cell transcriptome profiling of the human endometrium of patients with recurrent implantation failure. Theranostics. 2022;12(15):6527–47. Epub 2022/10/04. doi: 10.7150/thno.74053. PubMed PMID: 36185612; PubMed Central PMCID: PMCPMC9516226.

15. Zheng Y, Pan J, Xia C, Chen H, Zhou H, Ju W, et al. Characterization of placental and decidual cell development in early pregnancy loss by single-cell RNA sequencing. Cell Biosci. 2022;12(1):168. Epub 2022/10/09. doi: 10.1186/s13578-022-00904-5. PubMed PMID: 36209198; PubMed Central PMCID: PMCPMC9548121.

16. Yang Y, Zhu QY, Liu JL. Deciphering mouse uterine receptivity for embryo implantation at single-cell resolution. Cell Prolif. 2021;54(11):e13128. Epub 2021/09/25. doi: 10.1111/cpr.13128. PubMed PMID: 34558134; PubMed Central PMCID: PMCPMC8560620 as prejudicing the impartiality of the research reported.

17. Zheng ZH, Zhang GL, Jiang RF, Hong YQ, Zhang QY, He JP, et al. METTL3 is essential for normal progesterone signaling during embryo implantation via m(6)A-mediated translation control of progesterone receptor. Proc Natl Acad Sci U S A. 2023;120(5):e2214684120. Epub 2023/01/25. doi: 10.1073/pnas.2214684120. PubMed PMID: 36693099; PubMed Central PMCID: PMCPMC9945998.

18. Wu Y, Su K, Zhang Y, Liang L, Wang F, Chen S, et al. A spatiotemporal transcriptomic atlas of mouse placentation. Cell Discov. 2024;10(1):110. Epub 2024/10/23. doi: 10.1038/s41421-024-00740-6. PubMed PMID: 39438452; PubMed Central PMCID: PMCPMC11496649.

19. Xiao B, Wu S, Tian Y, Huang W, Chen G, Luo D, et al. Advances of NAT10 in diseases: insights from dual properties as protein and RNA acetyltransferase. Cell Biol Toxicol. 2024;41(1):17. Epub 2024/12/27. doi: 10.1007/s10565-024-09962-6. PubMed PMID: 39725720; PubMed Central PMCID: PMCPMC11671434.

20. Arango D, Sturgill D, Yang R, Kanai T, Bauer P, Roy J, et al. Direct epitranscriptomic regulation of mammalian translation initiation through N4-acetylcytidine. Mol Cell. 2022;82(15):2797–814 e11. Epub 2022/06/10. doi: 10.1016/j.molcel.2022.05.016. PubMed PMID: 35679869; PubMed Central PMCID: PMCPMC9361928.

21. Zhao W, Zhou Y, Cui Q, Zhou Y. PACES: prediction of N4-acetylcytidine (ac4C) modification sites in mRNA. Sci Rep. 2019;9(1):11112. Epub 2019/08/02. doi: 10.1038/s41598-019-47594-7. PubMed PMID: 31366994; PubMed Central PMCID: PMCPMC6668381.

22. Vasquez YM, Wang X, Wetendorf M, Franco HL, Mo Q, Wang T, et al. FOXO1 regulates uterine epithelial integrity and progesterone receptor expression critical for embryo implantation. PLoS Genet. 2018;14(11):e1007787. Epub 2018/11/20. doi: 10.1371/journal.pgen.1007787. PubMed PMID: 30452456; PubMed Central PMCID: PMCPMC6277115.

23. Soyal SM, Mukherjee A, Lee KY, Li J, Li H, DeMayo FJ, et al. Cre-mediated recombination in cell lineages that express the progesterone receptor. Genesis. 2005;41(2):58–66. Epub 2005/02/01. doi: 10.1002/gene.20098. PubMed PMID: 15682389.

24. Larrieu D, Britton S, Demir M, Rodriguez R, Jackson SP. Chemical inhibition of NAT10 corrects defects of laminopathic cells. Science. 2014;344(6183):527-32. Epub 2014/05/03. doi: 10.1126/science.1252651. PubMed PMID: 24786082; PubMed Central PMCID: PMCPMC4246063.

25. Yang W, Li HY, Wu YF, Mi RJ, Liu WZ, Shen X, et al. ac4C acetylation of RUNX2 catalyzed by NAT10 spurs osteogenesis of BMSCs and prevents ovariectomy-induced bone loss. Mol Ther Nucleic Acids. 2021;26:135–47. Epub 2021/09/14. doi: 10.1016/j.omtn.2021.06.022. PubMed PMID: 34513300; PubMed Central PMCID: PMCPMC8413676.

26. Balmus G, Larrieu D, Barros AC, Collins C, Abrudan M, Demir M, et al. Targeting of NAT10 enhances healthspan in a mouse model of human accelerated aging syndrome. Nat Commun. 2018;9(1):1700. Epub 2018/04/29. doi: 10.1038/s41467-018-03770-3. PubMed PMID: 29703891; PubMed Central PMCID: PMCPMC5923383 include Remodelin. The remaining authors declare no competing interests.

27. Brosens JJ, Hayashi N, White JO. Progesterone receptor regulates decidual prolactin expression in differentiating human endometrial stromal cells. Endocrinology. 1999;140(10):4809–20. Epub 1999/09/28. doi: 10.1210/endo.140.10.7070. PubMed PMID: 10499541.

28. Retis-Resendiz AM, Cid-Cruz Y, Velazquez-Hernandez DM, Romero-Reyes J, Leon-Juarez M, Garcia-Gomez E, et al. cAMP regulates the progesterone receptor gene expression through the protein kinase A pathway during decidualization in human immortalized endometrial stromal cells. Steroids. 2024;203:109363. Epub 2024/01/06. doi: 10.1016/j.steroids.2024.109363. PubMed PMID: 38182066.

29. Lydon JP, DeMayo FJ, Funk CR, Mani SK, Hughes AR, Montgomery CA, Jr., et al. Mice lacking progesterone receptor exhibit pleiotropic reproductive abnormalities. Genes Dev. 1995;9(18):2266–78. Epub 1995/09/15. doi: 10.1101/gad.9.18.2266. PubMed PMID: 7557380.

30. Wetendorf M, Wu SP, Wang X, Creighton CJ, Wang T, Lanz RB, et al. Decreased epithelial progesterone receptor A at the window of receptivity is required for preparation of the endometrium for embryo attachment. Biol Reprod. 2017;96(2):313–26. Epub 2017/02/17. doi: 10.1095/biolreprod.116.144410. PubMed PMID: 28203817; PubMed Central PMCID: PMCPMC6225975.

31. Shindoh H, Okada H, Tsuzuki T, Nishigaki A, Kanzaki H. Requirement of heart and neural crest derivatives-expressed transcript 2 during decidualization of human endometrial stromal cells in vitro. Fertil Steril. 2014;101(6):1781–90 e1-5. Epub 2014/04/22. doi: 10.1016/j.fertnstert.2014.03.013. PubMed PMID: 24745730.

32. Lynch VJ, Brayer K, Gellersen B, Wagner GP. HoxA-11 and FOXO1A cooperate to regulate decidual prolactin expression: towards inferring the core transcriptional regulators of decidual genes. PLoS One. 2009;4(9):e6845. Epub 2009/09/04. doi: 10.1371/journal.pone.0006845. PubMed PMID: 19727442; PubMed Central PMCID: PMCPMC2731163.

33. Liu JL, Wang TS. Systematic Analysis of the Molecular Mechanism Underlying Decidualization Using a Text Mining Approach. PLoS One. 2015;10(7):e0134585. Epub 2015/07/30. doi: 10.1371/journal.pone.0134585. PubMed PMID: 26222155; PubMed Central PMCID: PMCPMC4519252.

34. Gu Z, Zou L, Pan X, Yu Y, Liu Y, Zhang Z, et al. The role and mechanism of NAT10-mediated ac4C modification in tumor development and progression. MedComm (2020). 2024;5(12):e70026. Epub 2024/12/06. doi: 10.1002/mco2.70026. PubMed PMID: 39640362; PubMed Central PMCID: PMCPMC11617596.

35. Hewitt SC, Gruzdev A, Willson CJ, Wu SP, Lydon JP, Galjart N, et al. Chromatin architectural factor CTCF is essential for progesterone-dependent uterine maturation. FASEB J. 2023;37(8):e23103. Epub 2023/07/25. doi: 10.1096/fj.202300862R. PubMed PMID: 37489832; PubMed Central PMCID: PMCPMC10372848.

36. Wang T, Zou Y, Huang N, Teng J, Chen J. CCDC84 Acetylation Oscillation Regulates Centrosome Duplication by Modulating HsSAS-6 Degradation. Cell Rep. 2019;29(7):2078–91 e5. Epub 2019/11/14. doi: 10.1016/j.celrep.2019.10.028. PubMed PMID: 31722219.

37. Zheng J, Tan Y, Liu X, Zhang C, Su K, Jiang Y, et al. NAT10 regulates mitotic cell fate by acetylating Eg5 to control bipolar spindle assembly and chromosome segregation. Cell Death Differ. 2022;29(4):846–60. Epub 2022/02/26. doi: 10.1038/s41418-021-00899-5. PubMed PMID: 35210604; PubMed Central PMCID: PMCPMC8989979.

38. Mulac-Jericevic B, Mullinax RA, DeMayo FJ, Lydon JP, Conneely OM. Subgroup of reproductive functions of progesterone mediated by progesterone receptor-B isoform. Science. 2000;289(5485):1751-4. Epub 2000/09/08. doi: 10.1126/science.289.5485.1751. PubMed PMID: 10976068.

39. Liu KE, Hartman M, Hartman A, Luo ZC, Mahutte N. The impact of a thin endometrial lining on fresh and frozen-thaw IVF outcomes: an analysis of over 40 000 embryo transfers. Hum Reprod. 2018;33(10):1883–8. Epub 2018/09/22. doi: 10.1093/humrep/dey281. PubMed PMID: 30239738; PubMed Central PMCID: PMCPMC6145412.

40. Kelleher AM, DeMayo FJ, Spencer TE. Uterine Glands: Developmental Biology and Functional Roles in Pregnancy. Endocr Rev. 2019;40(5):1424–45. Epub 2019/05/11. doi: 10.1210/er.2018-00281. PubMed PMID: 31074826; PubMed Central PMCID: PMCPMC6749889.

41. Liu JL, Wang TS, Zhao M. Genome-Wide Association Mapping for Female Infertility in Inbred Mice. G3 (Bethesda). 2016;6(9):2929-35. Epub 2016/07/28. doi: 10.1534/g3.116.031575. PubMed PMID: 27449513; PubMed Central PMCID: PMCPMC5015949.

42. Schneider CA, Rasband WS, Eliceiri KW. NIH Image to ImageJ: 25 years of image analysis. Nat Methods. 2012;9(7):671–5. Epub 2012/08/30. doi: 10.1038/nmeth.2089. PubMed PMID: 22930834; PubMed Central PMCID: PMCPMC5554542.

43. Kim D, Paggi JM, Park C, Bennett C, Salzberg SL. Graph-based genome alignment and genotyping with HISAT2 and HISAT-genotype. Nat Biotechnol. 2019;37(8):907–15. Epub 2019/08/04. doi: 10.1038/s41587-019-0201-4. PubMed PMID: 31375807; PubMed Central PMCID: PMCPMC7605509.

44. Trapnell C, Williams BA, Pertea G, Mortazavi A, Kwan G, van Baren MJ, et al. Transcript assembly and quantification by RNA-Seq reveals unannotated transcripts and isoform switching during cell differentiation. Nat Biotechnol. 2010;28(5):511–5. Epub 2010/05/04. doi: 10.1038/nbt.1621. PubMed PMID: 20436464; PubMed Central PMCID: PMCPMC3146043.

45. Huang DW, Sherman BT, Tan Q, Kir J, Liu D, Bryant D, et al. DAVID Bioinformatics Resources: expanded annotation database and novel algorithms to better extract biology from large gene lists. Nucleic Acids Res. 2007;35(Web Server issue):W169-75. Epub 2007/06/20. doi: 10.1093/nar/gkm415. PubMed PMID: 17576678; PubMed Central PMCID: PMCPMC1933169.

46. Thorvaldsdottir H, Robinson JT, Mesirov JP. Integrative Genomics Viewer (IGV): high-performance genomics data visualization and exploration. Brief Bioinform. 2013;14(2):178–92. Epub 2012/04/21. doi: 10.1093/bib/bbs017. PubMed PMID: 22517427; PubMed Central PMCID: PMCPMC3603213.

47. Feng J, Liu T, Qin B, Zhang Y, Liu XS. Identifying ChIP-seq enrichment using MACS. Nat Protoc. 2012;7(9):1728–40. Epub 2012/09/01. doi: 10.1038/nprot.2012.101. PubMed PMID: 22936215; PubMed Central PMCID: PMCPMC3868217.

48. Cui X, Wei Z, Zhang L, Liu H, Sun L, Zhang SW, et al. Guitar: An R/Bioconductor Package for Gene Annotation Guided Transcriptomic Analysis of RNA-Related Genomic Features. Biomed Res Int. 2016;2016:8367534. Epub 2016/05/31. doi: 10.1155/2016/8367534. PubMed PMID: 27239475; PubMed Central PMCID: PMCPMC4864564.

49. Heinz S, Benner C, Spann N, Bertolino E, Lin YC, Laslo P, et al. Simple combinations of lineage-determining transcription factors prime cis-regulatory elements required for macrophage and B cell identities. Mol Cell. 2010;38(4):576–89. Epub 2010/06/02. doi: 10.1016/j.molcel.2010.05.004. PubMed PMID: 20513432; PubMed Central PMCID: PMCPMC2898526.

