## Supplementary figures and tables for "NAT10 governs uterine function and fertility by stabilizing progesterone receptor mRNA via ac^4^C modification"

### Supplementary figures and legends

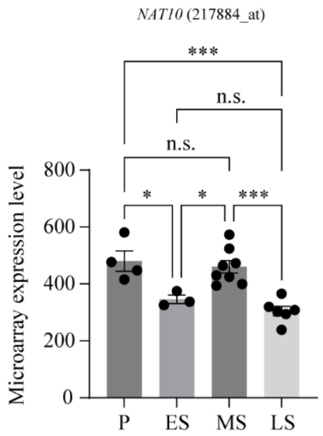


**Supplementary Figure 1. Expression of *NAT10* mRNA in human endometrium during the menstrual cycle in a microarray dataset GSE4888.** P, proliferative phase; ES, early secretory phase; MS, middle secretory phase; LS, late secretory phase. Data are presented as means ± SEM. n.s., not significant. *, $P < 0.05$; ***, $P < 0.001$.

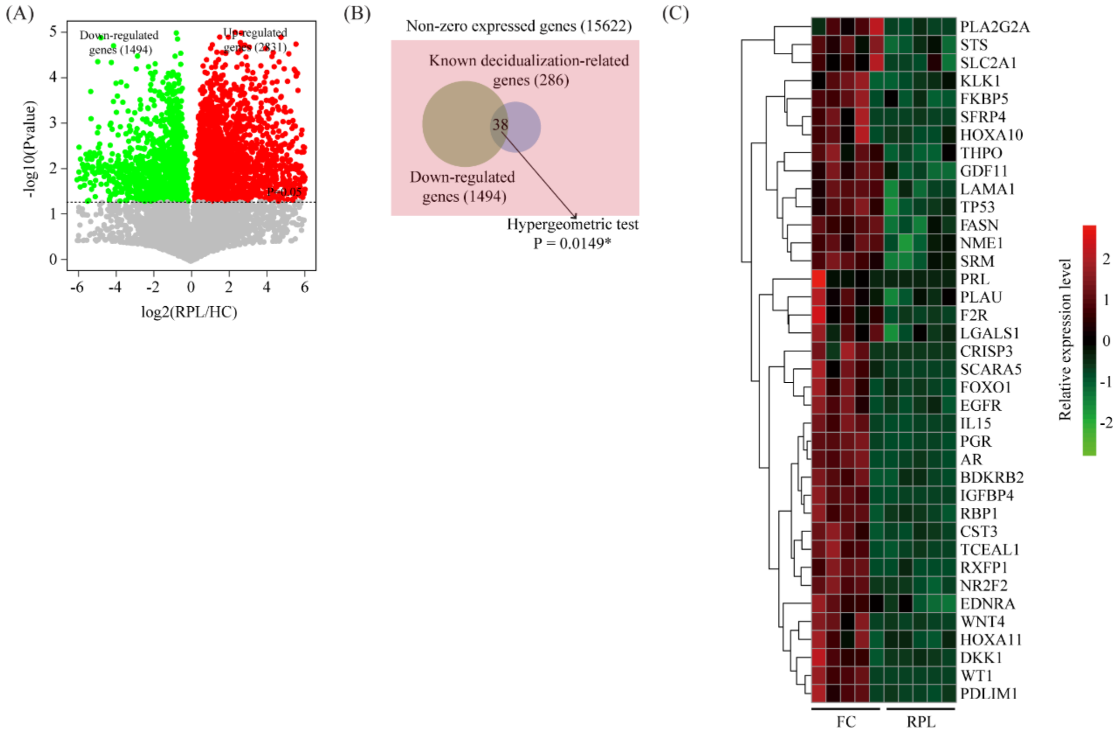


**Supplementary Figure 2. Down-regulated genes in hESCs from patients with RPL are enriched in known genes related to the decidualization process.** (A) Volcano plot showing significantly expressed genes in hESCs from RPL patients compared to FC patients based on GSE174399. Differentially expressed genes were identified using a t-test or Wilcoxon rank-sum test with a significance threshold of $P < 0.05$. (B) Venn diagram depicting the overlap of down-regulated genes and known decidualization-related genes, sourced from PMID: 26222155. (C) Heatmap plot showing the expression pattern of 38 overlap genes.

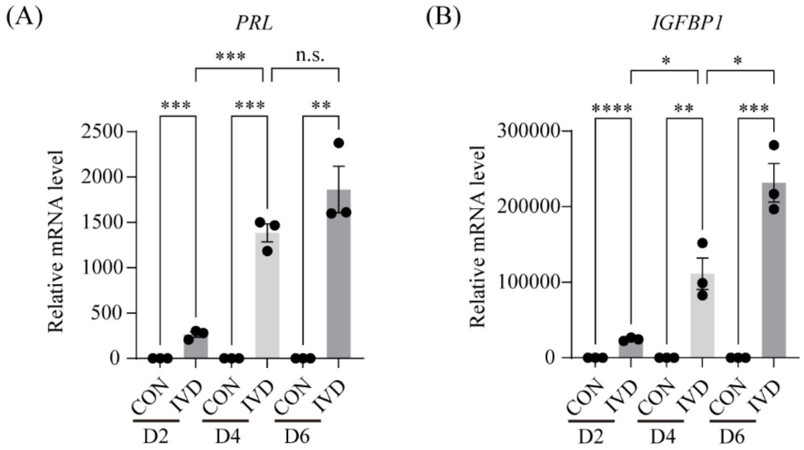


**Supplementary Figure 3. Decidualization marker PRL and IGFBP1 are highly induced in hESCs during in vitro decidualization.** Quantitative RT-PCR analysis of *PRL* (A) and *IGFBP1* (B) in primary hESCs during in vitro decidualization. CON, vehicle control; IVD, in vitro decidualization. Data are presented as mean ± SEM. *, P < 0.05; **, P < 0.01; ***, P < 0.001; ****, P < 0.0001.

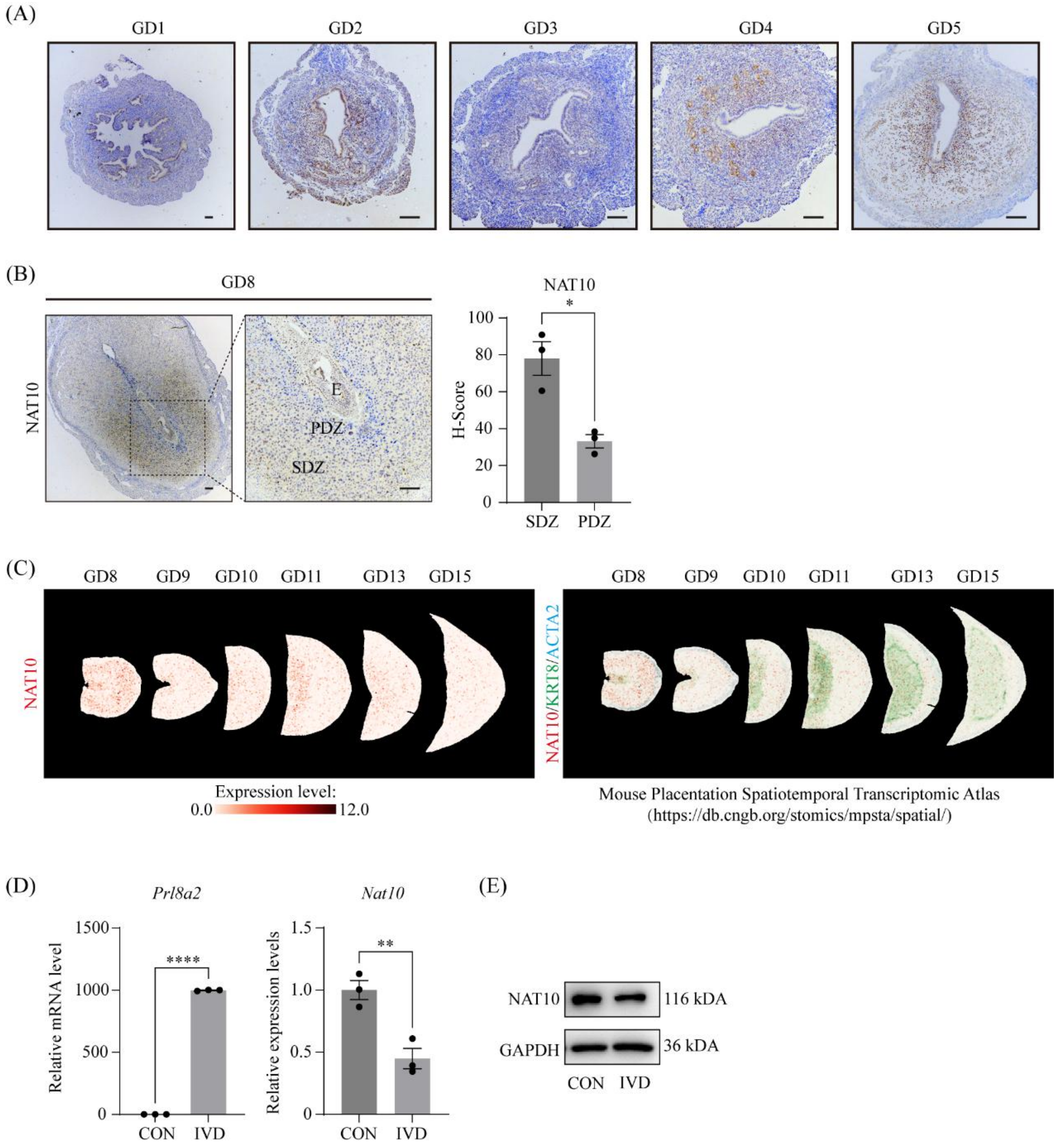


**Supplementary Figure 4. Expression of NAT10 in mouse uterus during pregnancy.** (A) Immunohistochemical staining for NAT10 protein in mouse uterus during the peri-implantation period. The brown color indicates positive staining for NAT10. GD, gestational day, with day 1 corresponding to the day of plug detection. Bar=100 µm. (B) Immunohistochemical staining for NAT10 protein in mouse uterus at the implantation sites on GD8. PDZ, primary decidual zone; SDZ, secondary decidual zone. Data are presented as mean ± SEM. *, $P < 0.05$ (C) Expression of NAT10 in mesometrial decidua or decidua basalis from GD8 to GD15 based on a Stereo-seq dataset (https://db.cngb.org/stomics/mpsta/spatial/). (D-E) Expression of NAT10 mRNA and protein in primary mouse endometrial stromal cells (mESCs) following in vitro decidualization for 3 days. Data are presented as mean ± SEM. **, $P < 0.01$; ****, $P < 0.0001$.

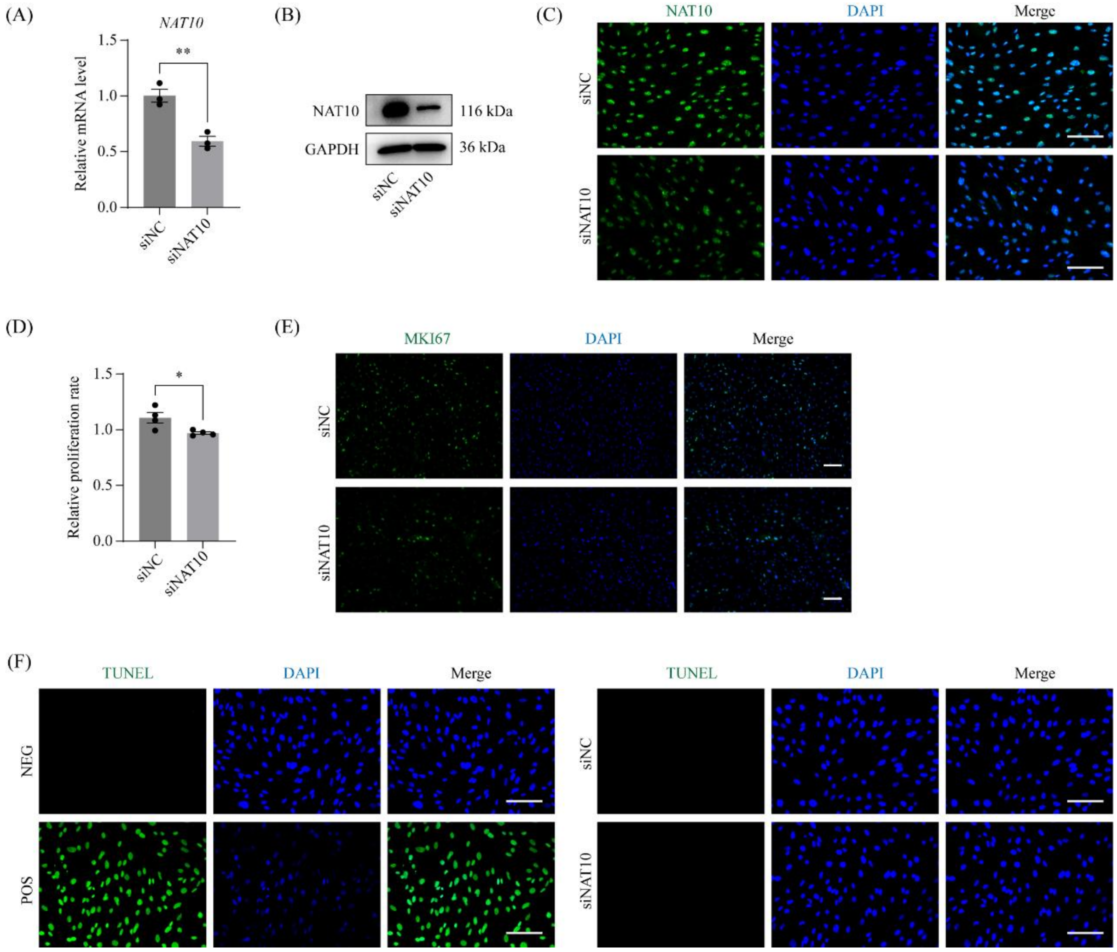


**Supplementary Figure 5. Knockdown of NAT10 in hESCs slows down cell proliferation without affecting apoptosis.** (A-C) Validation of NAT10-targeting siRNA in hESCs. (A) RNA levels of NAT10 after siRNA-mediated knockdown for 48h. Data are presented as mean ± SEM. **, $P < 0.01$. (B) Western blot analysis of NAT10 after siRNA-mediated knockdown for 48h. (C) Immunofluorescent staining of NAT10 protein in hESCs after siRNA-mediated knockdown for 48h. Bar = 100 µm. (D) Bar plot showing changes in cell proliferation rate after siRNA-mediated knockdown for 48h using the CCK8 kit. Data are presented as mean ± SEM. *, $P < 0.05$. (E) Immunofluorescent staining of MKI67 protein in hESCs after siRNA-mediated knockdown for 48h. Bar = 100 µm. (F) TUNEL staining to measure cell apoptosis after siRNA-mediated knockdown for 48h. NEG, no enzyme added to the TUNEL reaction as negative control; POS, DNase treatment before TUNEL reaction as positive control.

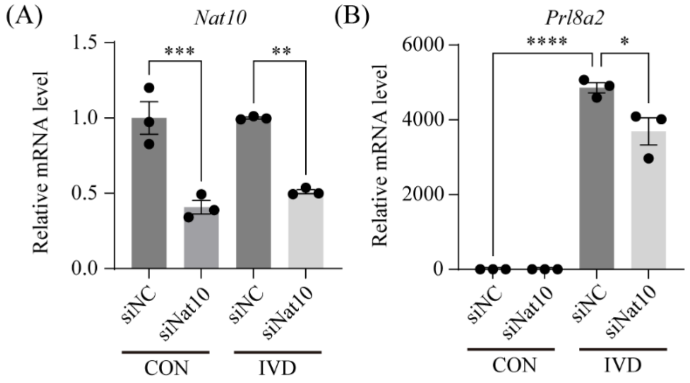


**Supplementary Figure 6. The expression of NAT10 in mouse endometrial stromal cells (mESCs) is required for in vitro decidualization.** Quantitative RT-PCR analysis of *Nat10* (A) and *Prl8a2* (B) mRNA expression in primary mESCs following Nat10 knockdown after in vitro decidualization for 3 days. CON, vehicle control; IVD, in vitro decidualization for 3 days. Data are presented as means ± SEM. **, P < 0.01; ***, P < 0.001.

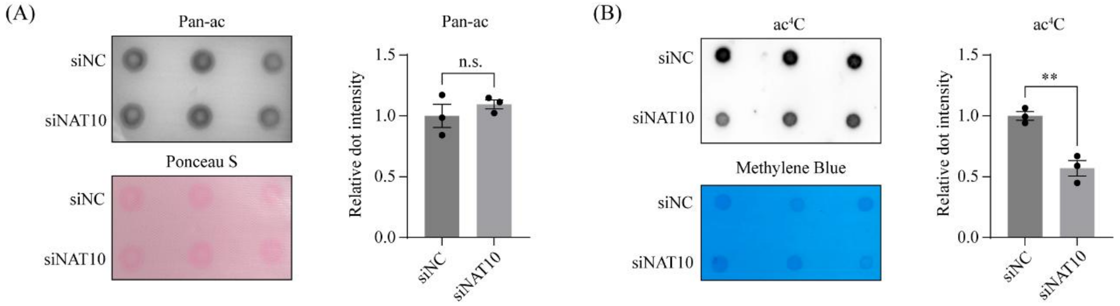


**Supplementary Figure 7. RNA acetylation is affected, but protein acetylation remains unchanged, in hESCs after NAT10 knockdown.** (A) Dot blot analysis of protein acetylation levels using an anti-pan-acetylation (Pan-ac) antibody in hESCs 48h after Nat10 knockdown. Data are presented as means ± SEM. n.s., not significant. (B) Dot blot analysis of RNA acetylation level using an anti-$ac^4C$ antibody in hESCs 48h after Nat10 knockdown. Data are presented as means ± SEM. **, P < 0.01.

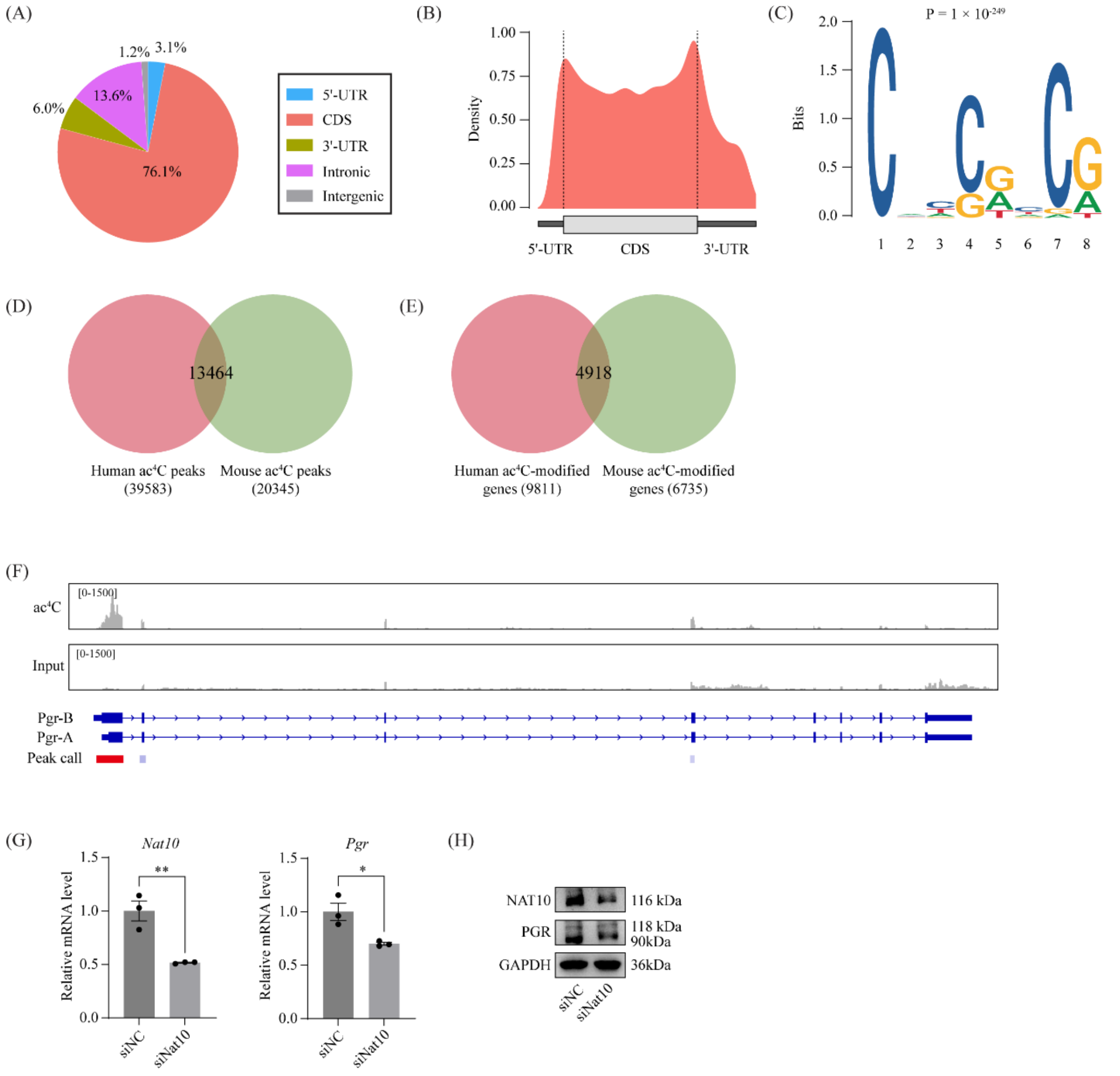


**Supplementary Figure 8. PGR is a highly conserved target of NAT10 in mice.** (A-C) Global profiling of ac$^4$C modification in mouse uterus on GD4 using acRIP-seq. (A) Pie chart presenting fractions of ac$^4$C peaks in different genomic segments. (B) The metagene distribution of ac$^4$C peaks in the gene body. (C) Sequence logo representing the consensus motif of ac$^4$C modification. (D-E) Venn diagram depicting the overlap of ac$^4$C modification between humans and mice at the peak (D) and the gene (E) levels, respectively. (F) Integrative genomics viewer (IGV) snapshot displaying the coverage of ac$^4$C immunoprecipitation and input control in mouse PGR mRNA. (G-H) Expression of NAT10 mRNA and protein in mESCs after 48h of NAT10 knockdown. Data are presented as mean ± SEM. *, $P < 0.05$; **, $P < 0.01$.

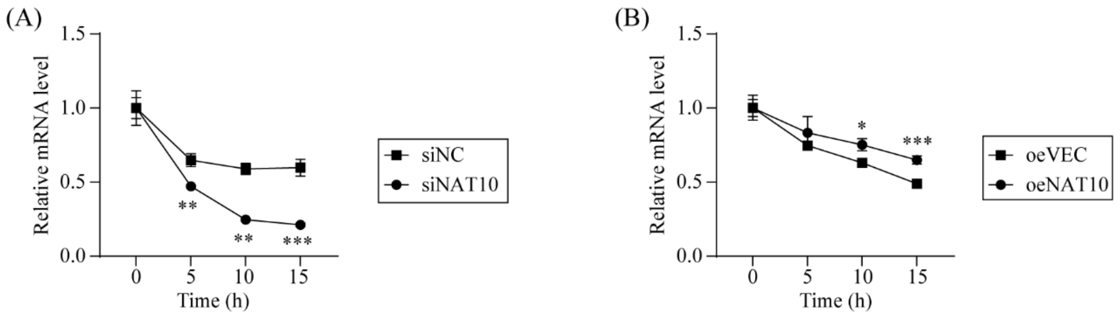


**Supplementary Figure 9. The stability of PGR mRNA is enhanced by NAT10.** (A) RNA stability assay showing the mRNA levels of PGR in hESCs following NAT10 knockdown. Cells were treated with actinomycin D 48h after NAT10 knockdown. Data are presented as mean ± SEM. **, P < 0.01; ***, P < 0.001. (B) RNA stability assay showing the mRNA levels of PGR in hESCs following NAT10 overexpression. Data are presented as mean ± SEM. *, P < 0.05; ***, P < 0.001.

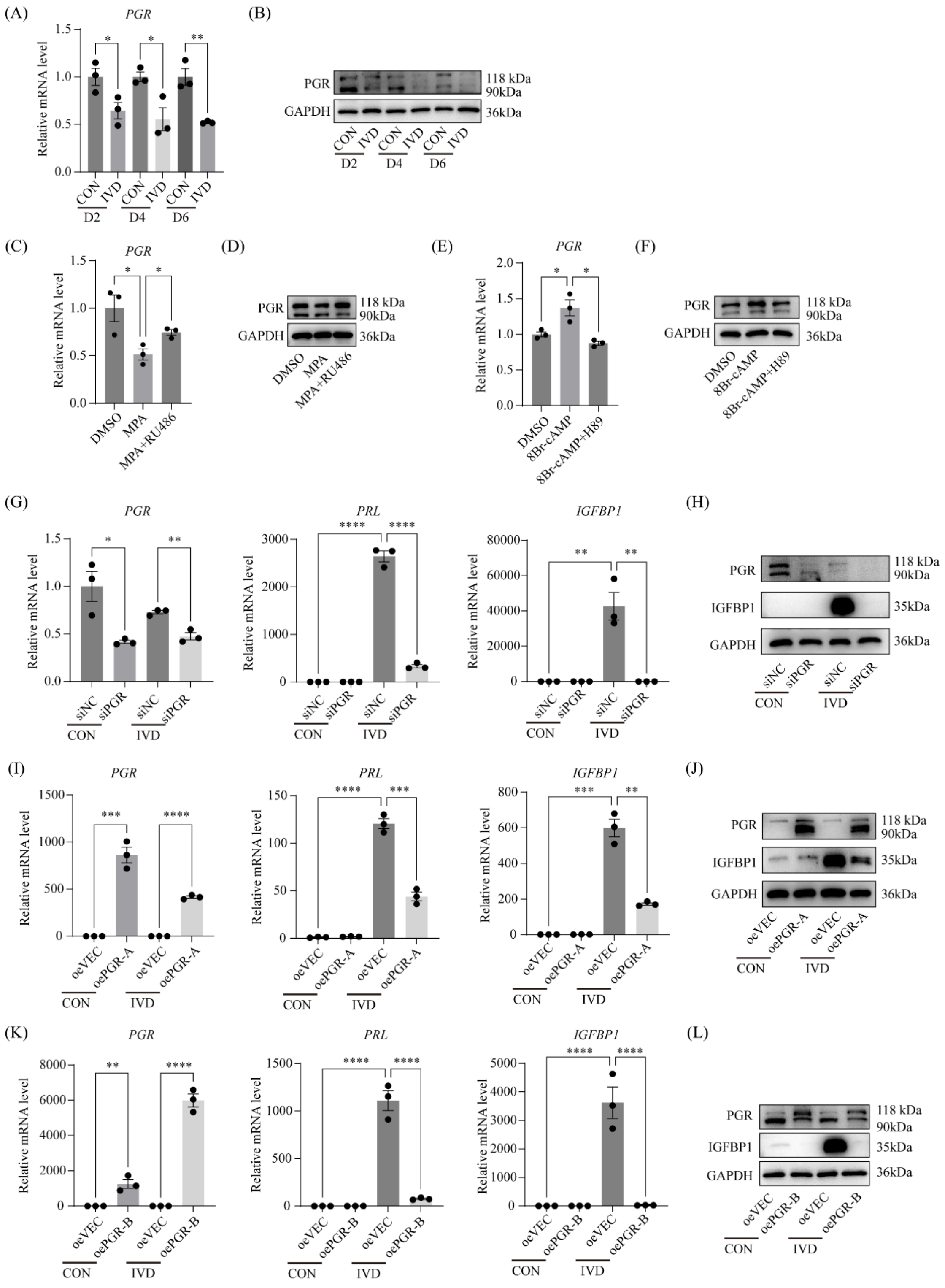


**Supplementary Figure 10. The expression, regulation and function of PGR in hESCs during in vitro decidualization.** (A-B) Expression of *PGR* mRNA and protein in decidualizing hESCs at different time points. CON, vehicle control; IVD, in vitro decidualization. Data are presented as mean ± SEM. *, P < 0.05; **, P < 0.01. (C-D) Expression of PGR mRNA and protein in hESCs treated MPA or MPA plus RU486 for 48h. Data are presented as mean ± SEM. *, P < 0.05. (E-F) Expression of PGR mRNA and protein in hESCs treated with 8Br-cAMP or 8Br-cAMP plus H89 for 48h. Data are presented as mean ± SEM. *, P < 0.05. (G) The mRNA expression levels of PRL and IGFBP1 in hESCs following PGR knockdown and subsequent decidualization for 4 days. Data are presented as mean ± SEM. *, P < 0.05; **, P < 0.01; ****, P < 0.0001. (H) Protein expression levels of IGFBP1 in hESCs following PGR knockdown and subsequent decidualization for 4 days. (I) Quantitative RT-PCR analysis of PRL and IGFBP1 mRNA expression in hESCs following PGR-A overexpression and subsequent decidualization for 4 days. Data are presented as means ± SEM. **, P < 0.01; ***, P < 0.001; ****, P < 0.0001. (J) IGFBP1 protein expression in hESCs following PGR-A overexpression and subsequent decidualization for 4 days. (K) Quantitative RT-PCR analysis of PRL and IGFBP1 mRNA expression in hESCs following PGR-B overexpression and subsequent decidualization for 4 days. Data are presented as means ± SEM. **, P < 0.01; ****, P < 0.0001. (L) IGFBP1 protein expression in hESCs following PGR-B overexpression and subsequent decidualization for 4 days.

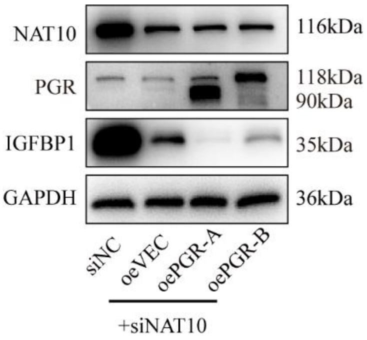


**Supplementary Figure 11. Overexpression of neither PGR-A nor PGR-B is able to restore decidualization in hESCs with NAT10 knockdown.** Cells were harvested at day 4 of decidualization.

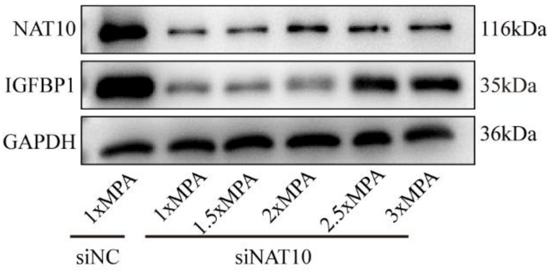


**Supplementary Figure 12. Screening for an optimal concentration of MPA to rescue decidualization in hESCs with NAT10 knockdown.** Cells were harvested at day 4 of decidualization. Notably, the concentrations of 2.5-fold and 3-fold MPA were effective in restoring decidualization, with the 3-fold concentration showing slightly better results compared to the 2.5-fold concentration. Therefore, the 3-fold concentration of MPA was selected for further experiments.

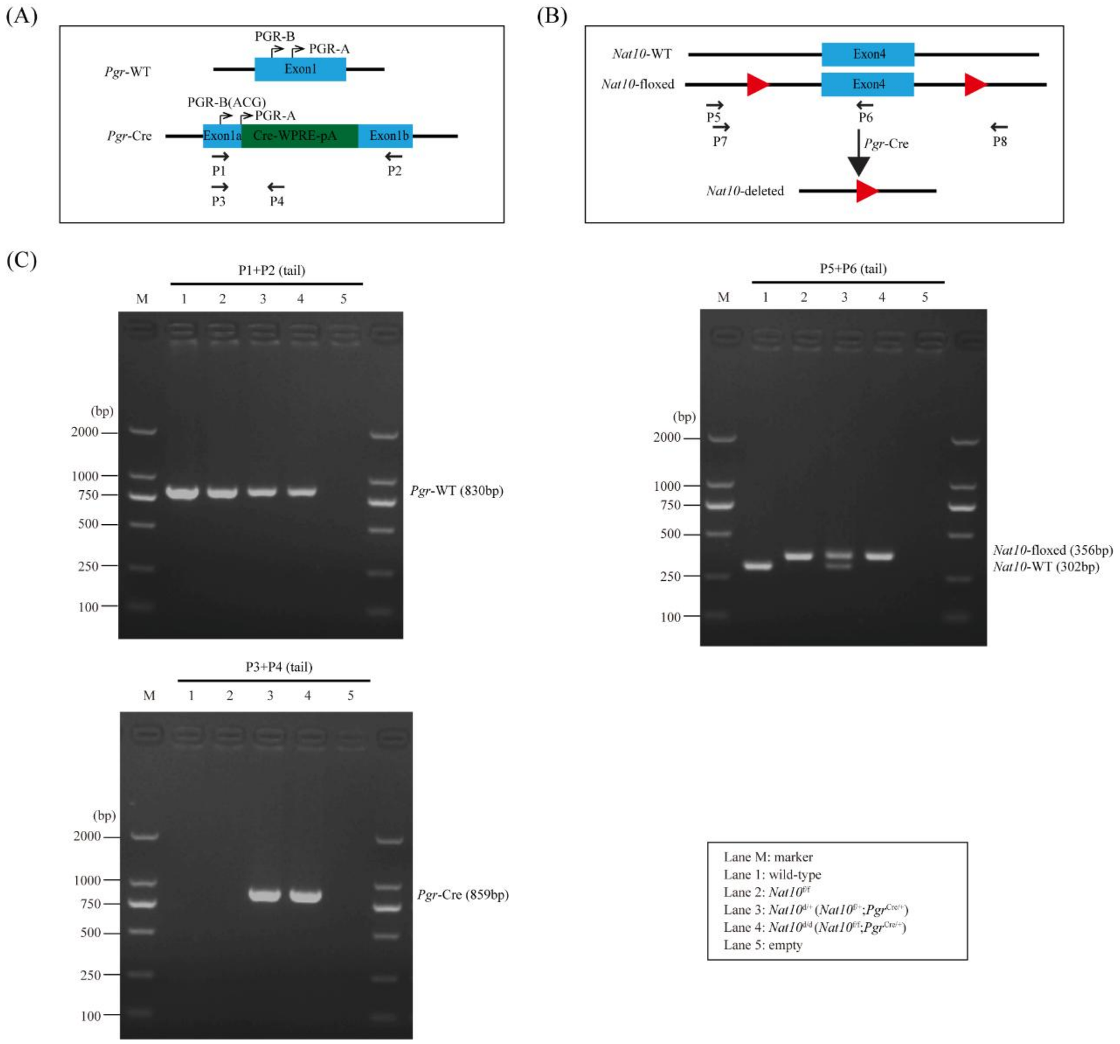


**Supplementary Figure 13. Genotyping analysis of *Nat10*$^{d/d}$ mice.** (A) Diagram showing genotyping primers for the *Pgr*-Cre allele. (B) Diagram showing genotyping primers for the *Nat10*-floxed allele. P7/P8 are used in Fig. 4B. (C) Genotyping PCR analysis for *Nat10*$^{d/d}$ and *Nat10*$^{f/f}$ mice.

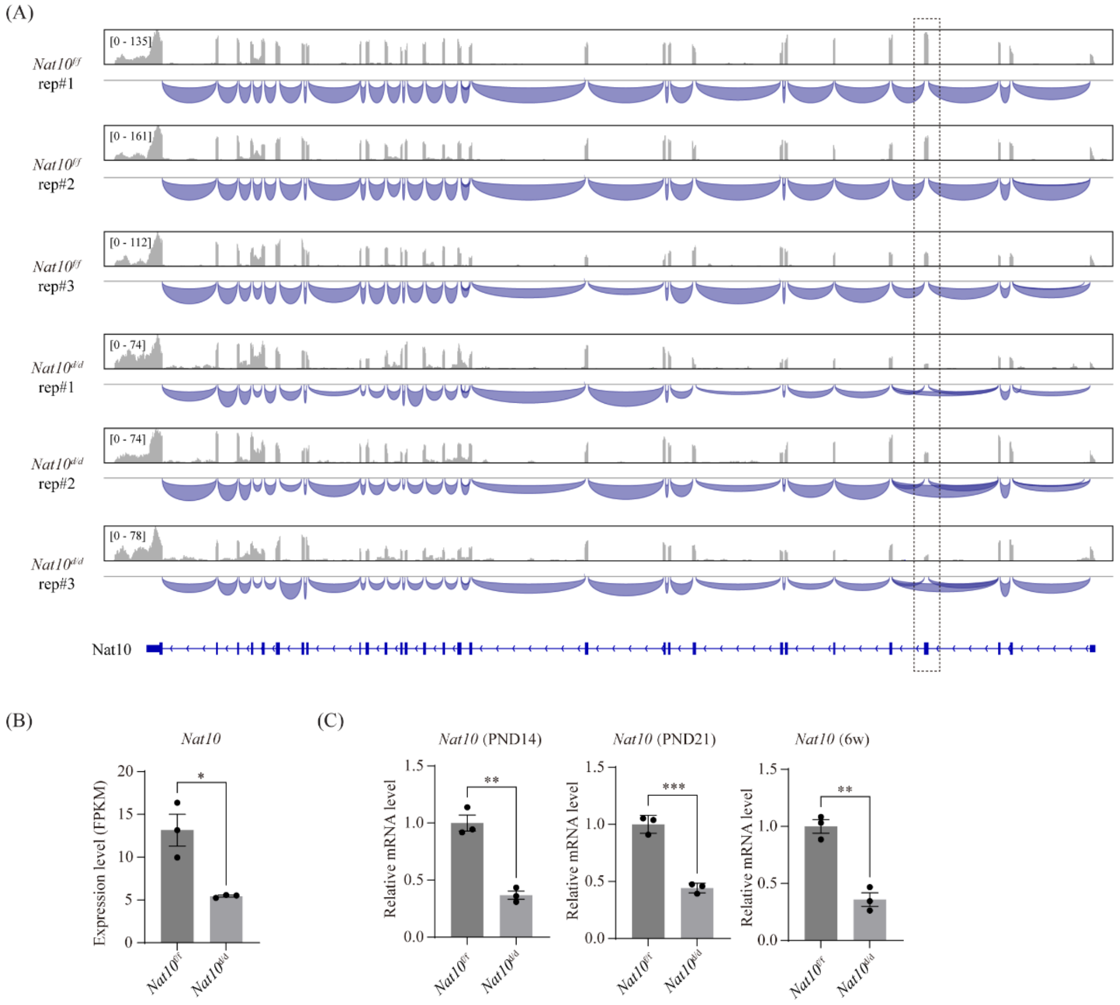


**Supplementary Figure 14. Deletion of exon 4 in *Nat10*$^{d/d}$ mouse uterus causes alternative splicing and reduced mRNA levels.** (A) Integrative Genomics Viewer showing the deletion of exon 4 of *Nat10* in the uterus of *Nat10*$^{d/d}$ mice at PND21 based on RNA-seq analysis. (B) The mRNA expression levels of *Nat10* in the uterus from *Nat10*$^{d/d}$ and *Nat10*$^{f/f}$ mice based on RNA-seq data. Data are presented as mean ± SEM. *, $P < 0.05$. (C) Validation of Nat10 mRNA expression levels in the uterus from *Nat10*$^{d/d}$ and *Nat10*$^{f/f}$ mice by using quantitative RT-PCR. Data are presented as mean ± SEM. **, $P < 0.01$; ***, $P < 0.001$.

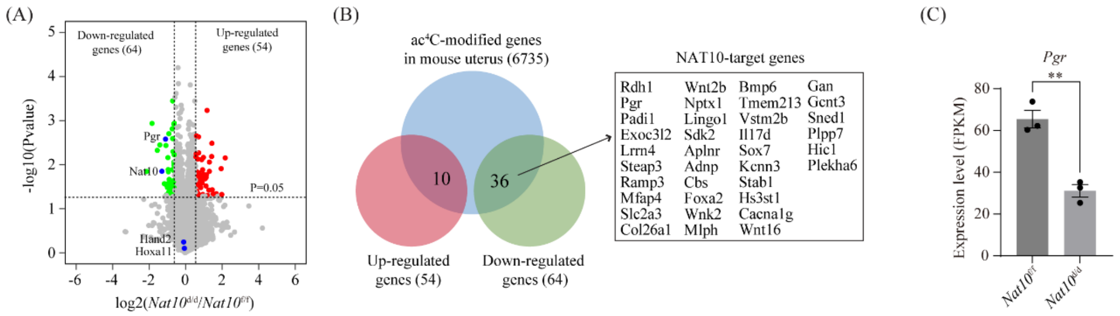


**Supplementary Figure 15. RNA-seq analysis reveals that *Pgr* mRNA was down-regulated in *Nat10*$^{d/d}$ uterus on PND21.** (A) Volcano plot for differentially expressed genes in the uterus from *Nat10*$^{d/d}$ mice and *Nat10*$^{f/f}$ mice on PND21, based on RNA-seq analysis. (B) Venn diagram depicting the overlap of differentially expressed genes identified by RNA-seq and mouse ac$^4$C-modified genes identified by acRIP-seq. (C) Bar plot showing reduced *Pgr* mRNA in the uterus of *Nat10*$^{d/d}$ mice compared to Nat10$^{f/f}$ mice, based on RNA-seq data. **, P < 0.01.

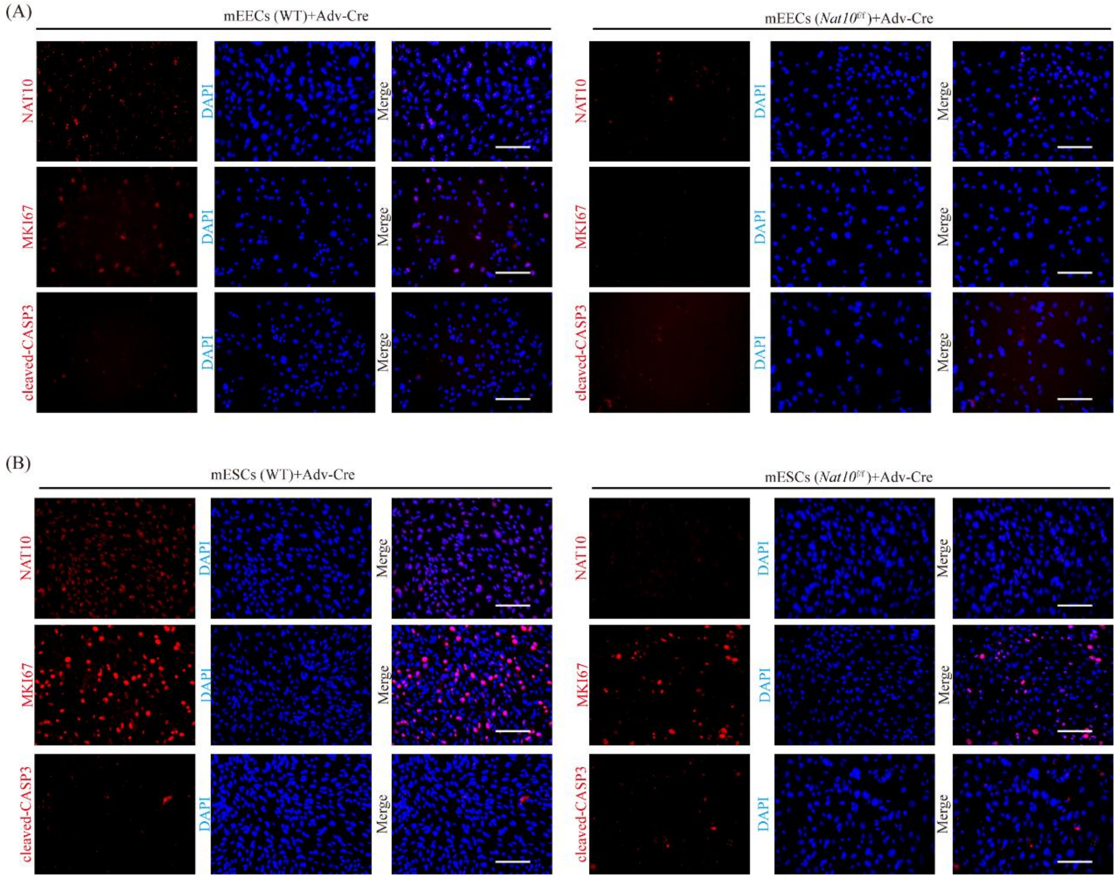


**Supplementary Figure 16. Evaluation of the consequences of *Nat10* deletion in mEECs and mESCs in vitro by Adv-Cre.** (A) Immunofluorescent staining of NAT10, MKI67 and cleaved-CASP3 in cultured mouse endometrial epithelial cells (mEECs) from *Nat10*$^{f/f}$ mice (WT mice serving as control) following treatment with Adv-Cre for 4 days. Bar = 100 µm. (B) Immunofluorescent staining of NAT10, MKI67 and cleaved-CASP3 in cultured mouse stromal cells (mESCs) from *Nat10*$^{f/f}$ mice (WT mice serving as control) following treatment with Adv-Cre for 4 days. Bar = 100 µm.

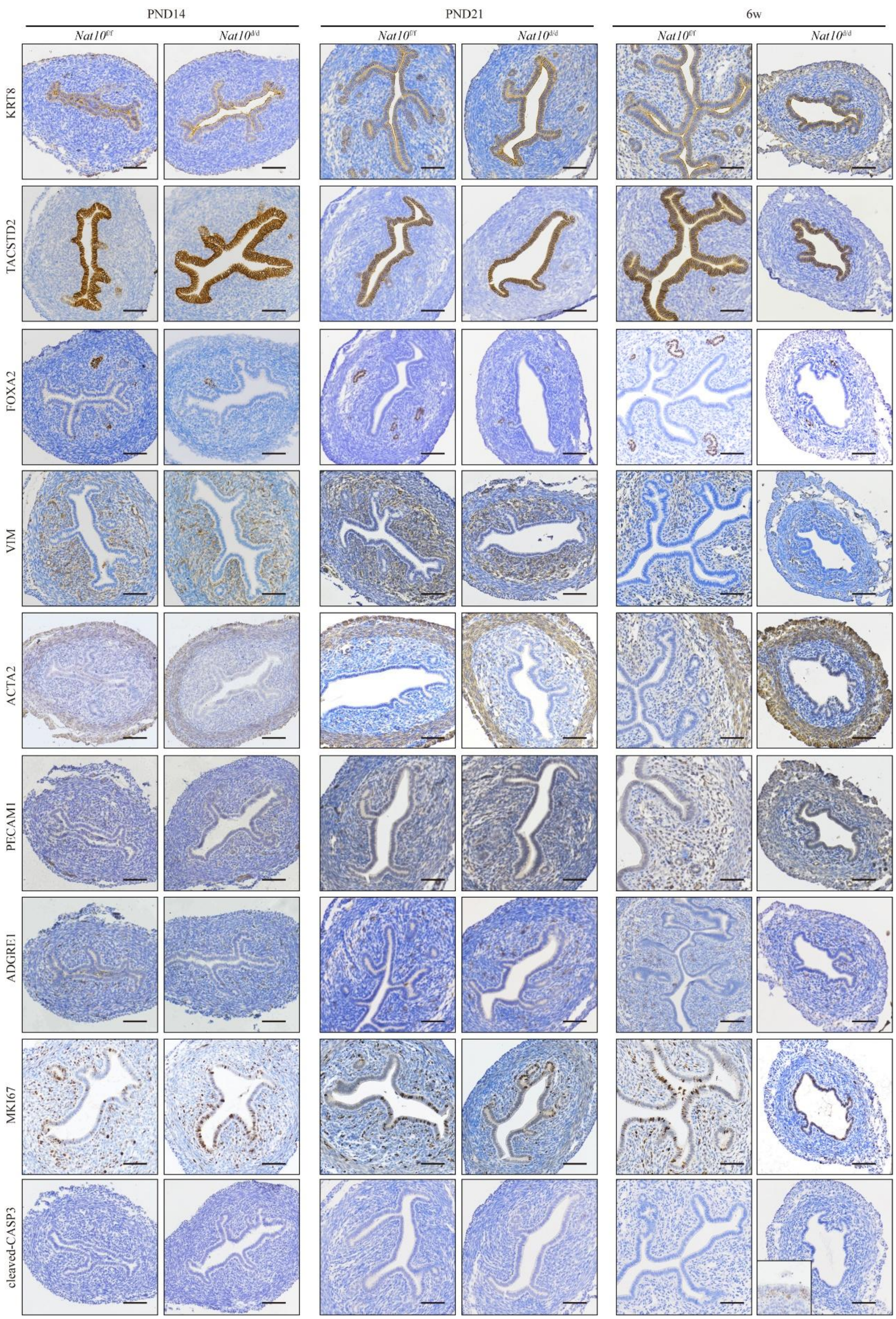


**Supplementary Figure 17. Examination of structural alterations in the *Nat10*-deleted uterus during pubertal development through immunohistochemical staining of cell-type specific markers.** For cleaved-CASP3, luminal epithelial cells are used as positive control (shown as inset). Bar = 50 μm.

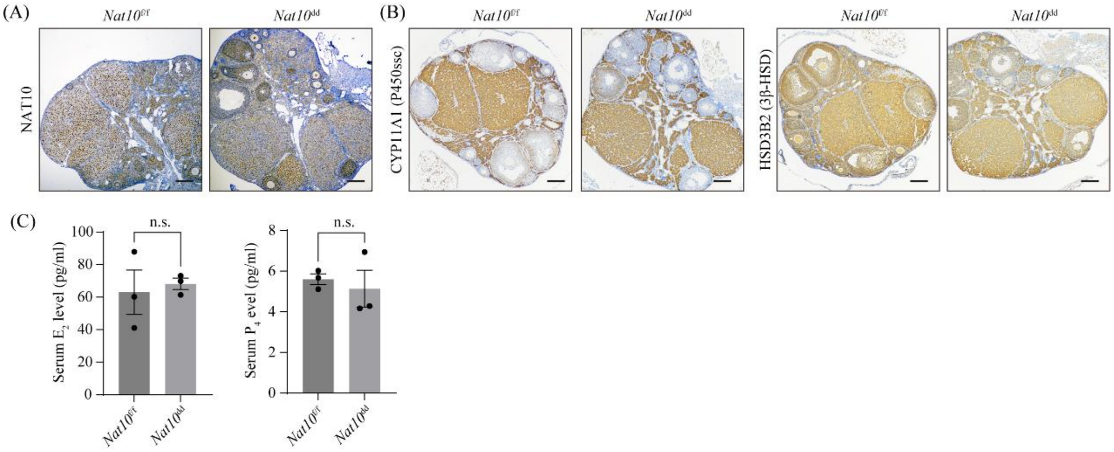


**Supplementary Figure 18. Ovarian function in conditional Nat10 deletion mice appears to be normal.** (A) Immunohistochemical staining of NAT10 in the ovary of *Nat10*$^{d/d}$ and *Nat10*$^{f/f}$ mice on GD4. Bar = 100 µm. (B) Immunohistochemical staining of CYP11A1 and HSD3B2 in the ovary of *Nat10*$^{d/d}$ and *Nat10*$^{f/f}$ mice on GD4. Bar = 100 µm. (C) The concentration of circulating $E_2$ and $P_4$ in *Nat10*$^{d/d}$ mice and *Nat10*$^{f/f}$ mice on GD4. The concentrations were measured using ELISA kits (Genkern, Guangzhou, China), according to the manufacturer's instructions. n.s., not significant.

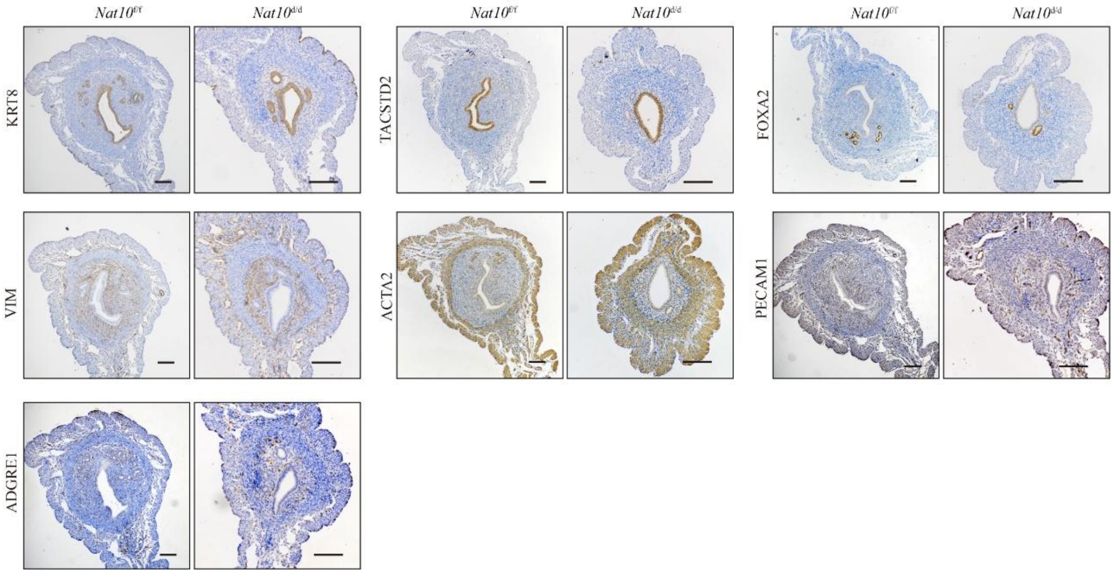


**Supplementary Figure 19. Examination of structural alterations in the *Nat10*-deleted uterus on GD4 through immunohistochemical staining of cell-type specific markers.** Bar = 100 µm.

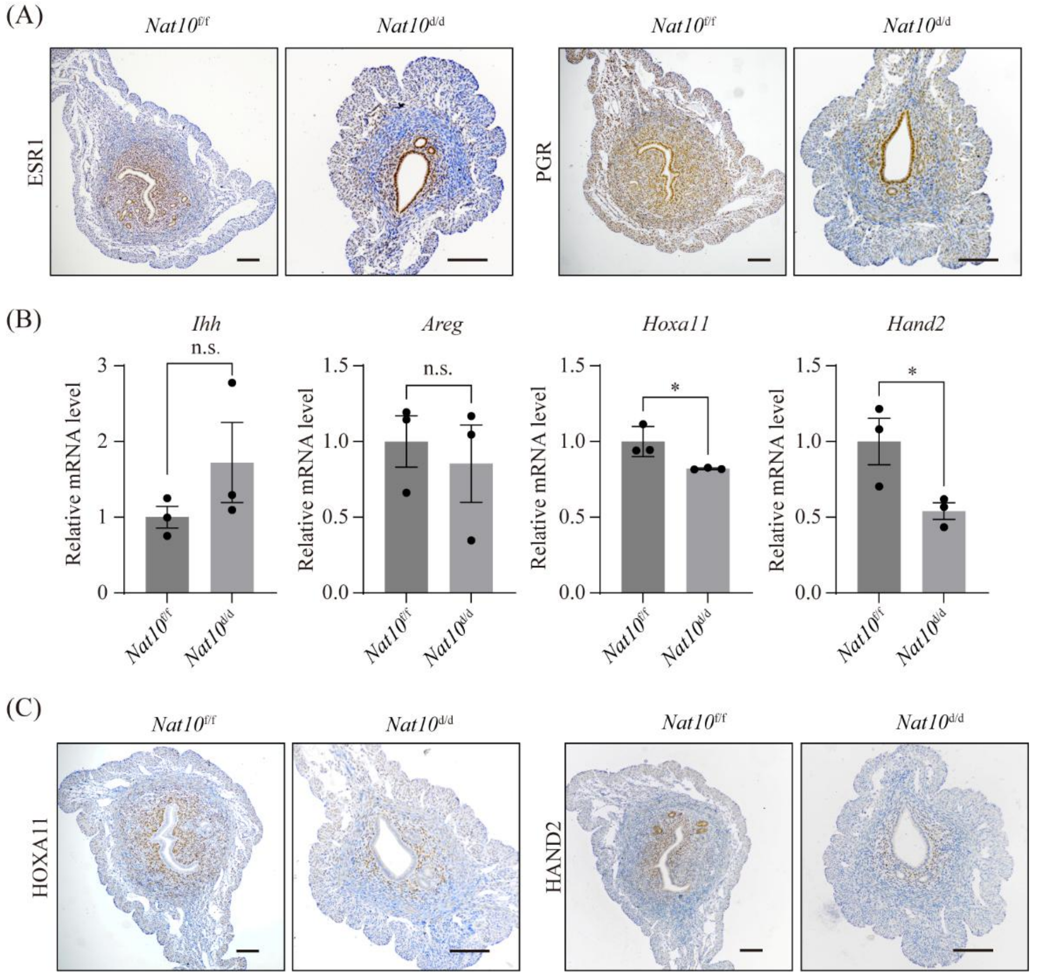


**Supplementary Figure 20. Deletion of Nat10 leads to decreased expression of PGR and its stromal targets HOXA11 and HAND2 in the uterus on GD4.** (A) Immunohistochemistry staining of PGR and ESR1 in the uterus from *Nat10*$^{f/f}$ mice and *Nat10*$^{d/d}$ mice on GD4. Bar=100 µm. (B) Quantitative RT-PCR analysis of $P_4$ taget genes in the uterus from *Nat10*$^{f/f}$ and *Nat10*$^{d/d}$ mice on GD4. Data are presented as means ± SEM. *, P < 0.05. (C) Immunohistochemistry staining of HOXA11 and PGR in the uterus from *Nat10*$^{f/f}$ mice and *Nat10*$^{d/d}$ mice on GD4. Bar=100 µm.

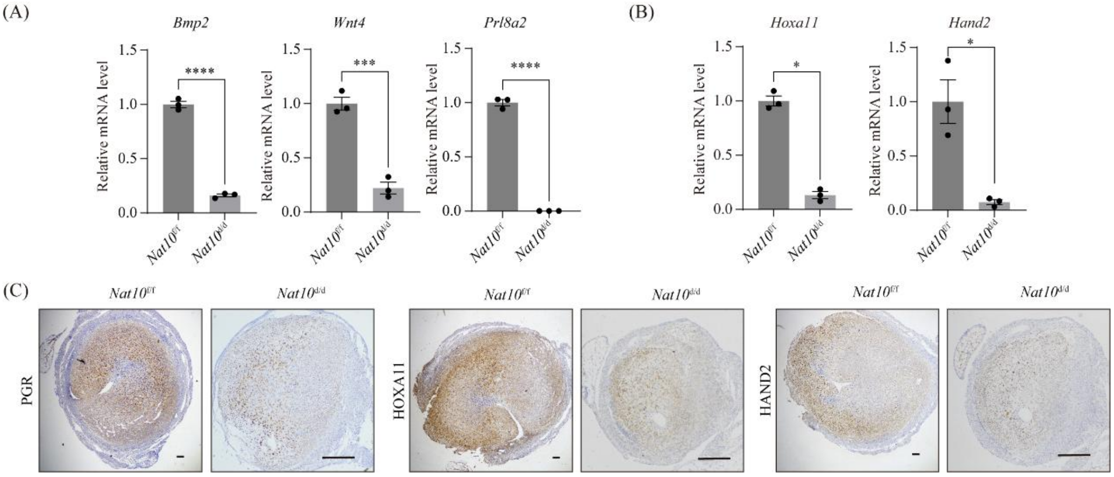


**Supplementary Figure 21. Decidualization failure in *Nat10*$^{d/d}$ mice following artificial decidualization is associated with the down-regulation of PGR, HOXA11 and HAND2.** (A) Quantitative RT-PCR analysis of decidualization marker genes in the stimulated uterine horn of *Nat10*$^{d/d}$ mice and *Nat10*$^{f/f}$ mice following artificial decidualization. Data are presented as means ± SEM. ***, $P < 0.001$; ****, $P < 0.0001$. (B) Quantitative RT-PCR analysis of HOXA11 and HAND2 in the stimulated uterine horn of *Nat10*$^{d/d}$ mice and *Nat10*$^{f/f}$ mice. Data are presented as means ± SEM. *, $P < 0.05$. (C) Immunohistochemistry staining of NAT10, HOXA11 and HAND2 in the stimulated uterine horn from *Nat10*$^{d/d}$ mice and *Nat10*$^{f/f}$ mice. Bar=100 μm.

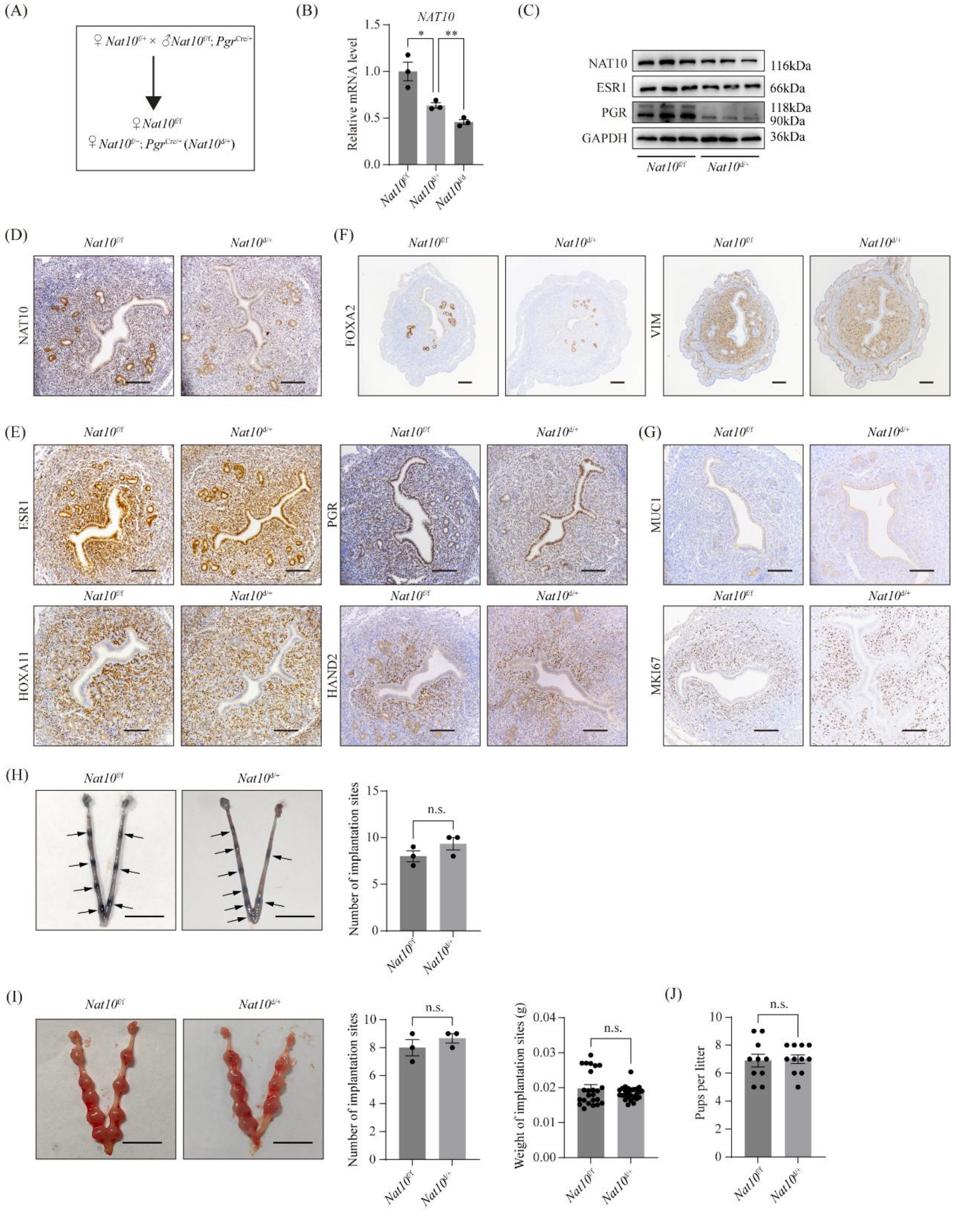

**Supplementary Figure 22. Heterozygous *Nat10* conditional knockout mice exhibit decreased NAT10 expression in the uterus, yet maintain normal fertility.** (A) A diagram of the breeding scheme for generating *Nat10*$^{d/+}$ mice. (B) Quantitative RT-PCR analysis of *Nat10* expression in the uterus on GD4 from *Nat10*$^{d/d}$, *Nat10*$^{d/+}$ and *Nat10*$^{f/f}$ mice. Data are presented as means ± SEM. *, P < 0.05; **, P < 0.01. (C) Western blot analysis of NAT10, ESR1 and PGR protein levels in the uterus of *Nat10*$^{d/+}$ mice and *Nat10*$^{f/f}$ mice on GD4. (D-G) Immunohistochemistry staining of NAT10, ESR1, PGR, HOXA11, HAND2, FOXA2, VIM, MUC1 and MKI67 in the uterus from *Nat10*$^{d/+}$ mice and *Nat10*$^{f/f}$ mice on GD4. Bar=100 μm. (H) Bar plot showing the number of embryo implantation sites in *Nat10*$^{d/+}$ mice and *Nat10*$^{f/f}$

mice on GD5. Implantation sites were visualized by the blue dye injection method. Implantation sites are marked by arrowheads. Bar = 1 cm. Data are presented as mean ± SEM. n.s., not significant. (I) Analysis of the number and the weight of embryo implantation sites in *Nat10*$^{d/+}$ mice and *Nat10*$^{f/f}$ mice on GD8. n.s., not significant. (J) Analysis of litter sizes for 3 *Nat10*$^{d/+}$ mice and 3 *Nat10*$^{f/f}$ mice during the 4-month fertility test. Data are presented as mean ± SEM. n.s., not significant.

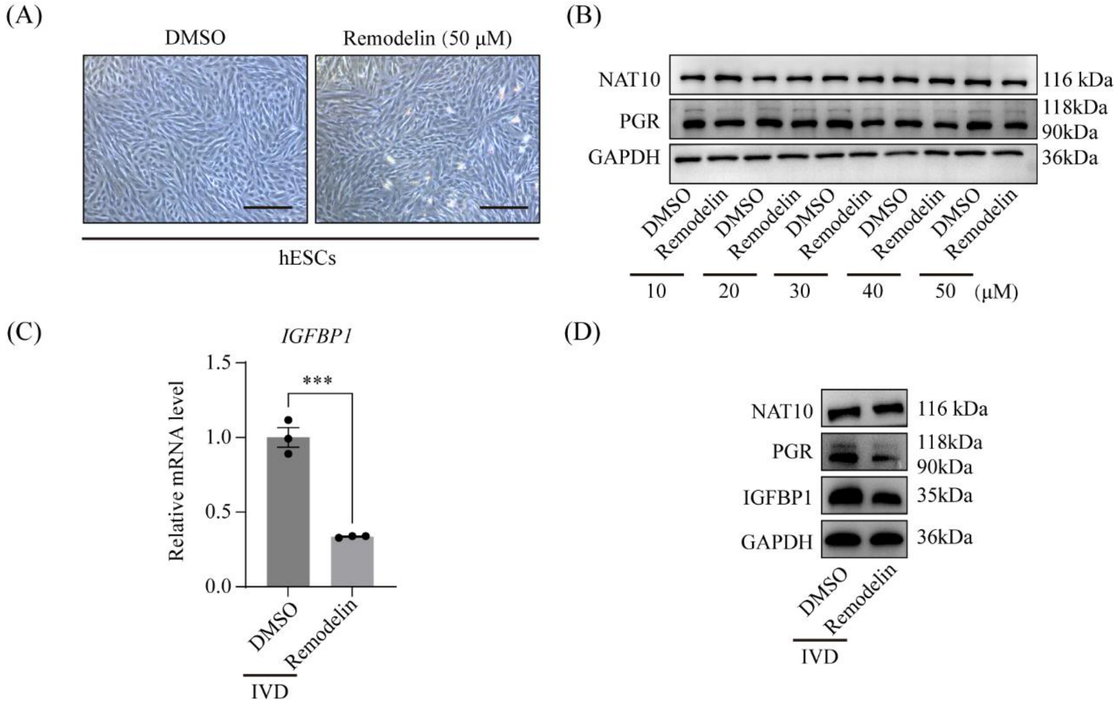


**Supplementary Figure 23. Remodelin treatment impairs hESC decidualization in vitro.** (A) The morphological appearance of the hESCs remains normal when treated with a concentration of Remodelin up to 50 μM for 48h. Bar = 100 μm. (B) Screening for an effective concentration of Remodelin in hESCs using PGR protein expression level as an indicator. Cells were harvested 48 hours post-treatment with Remodelin. Notably, a down-regulation of PGR protein was observed at concentrations of 30 μM and above. (C-D) The impact of Remodelin on hESC decidualization in vitro. (C) Quantitative RT-PCR analysis of IGFBP1 mRNA expression in hESCs following 30 μM Remodelin treatment in the in vitro decidualization model. CON, vehicle control; DEC, in vitro decidualization for 4 days. Data are presented as means ± SEM. ***, $P < 0.001$. (D) PGR and IGFBP1 protein expression in hESCs following 30 μM Remodelin treatment in the in vitro decidualization model.

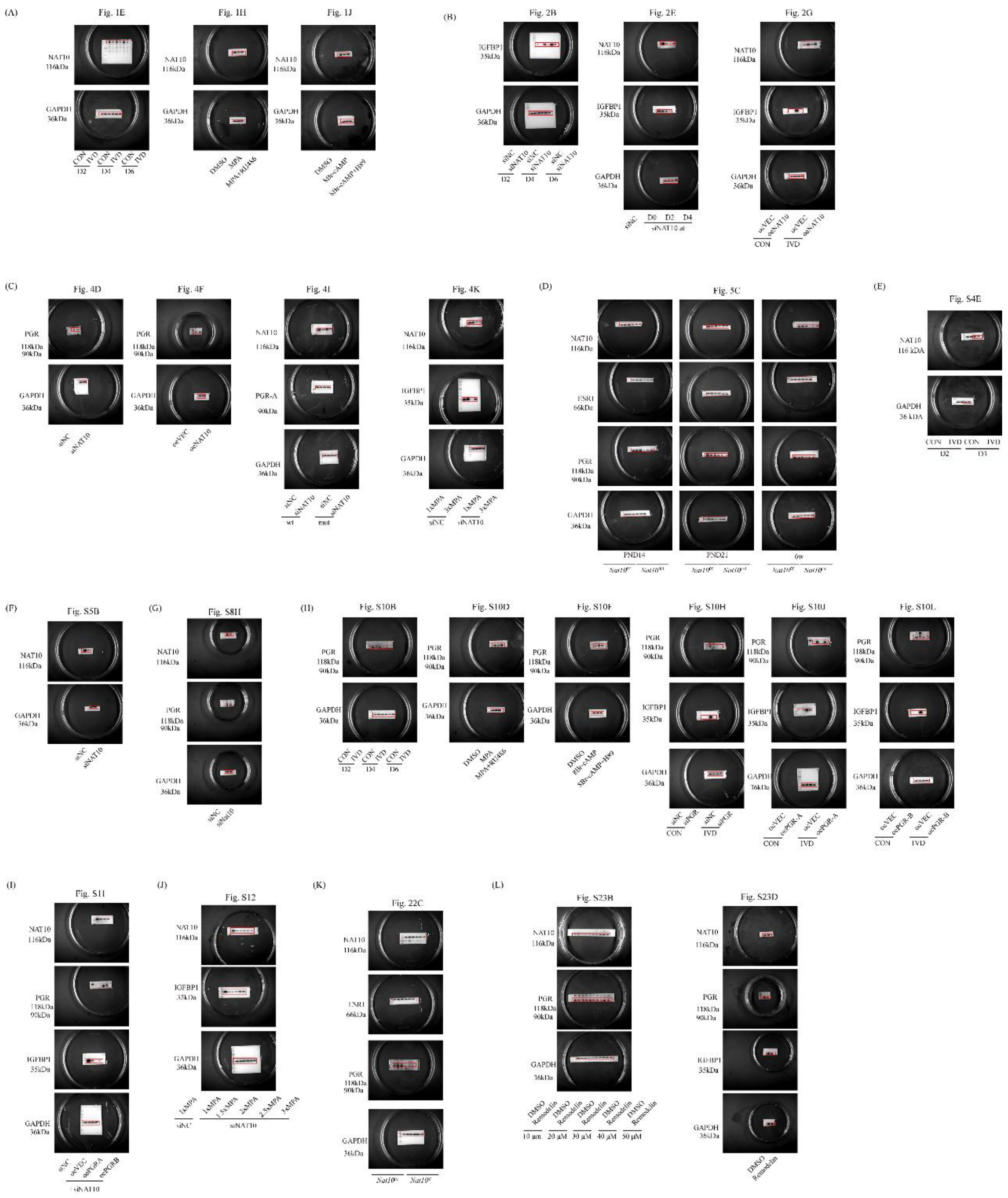

**Supplementary Figure 24. Pictures of the uncropped Western blots.**

**Supplementary tables**

**Supplementary Table 1. The complete list of $ac^4C$ peaks in human endometrial tissues at the middle secretory phase (q-value < 0.05).**

**Supplementary Table 2. Differentially expressed genes in NAT10 knockdown hESCs compared to control (fold change > 1.5 and p-value < 0.05).**

**Supplementary Table 3. The complete list of $ac^4C$ peaks in mouse uterus on GD4 (q-value < 0.05).**

**Supplementary Table 4. Differentially expressed genes in *Nat10*$^{d/d}$ uterus compared to *Nat10*$^{f/f}$ uterus at PND21 (fold change > 1.5 and p-value < 0.05).**

**Supplementary Table 5. Detailed information of human participants for isolation of primary endometrial stromal cells in this study.**

**Supplementary Table 6. Primers used in this study.**

**Supplementary Table 7. Antibodies used in this study.**

**Supplementary Table 8. siRNAs and overexpression vectors used in this study.**
